# Molecular basis for dyneinopathies reveals insight into dynein regulation and dysfunction

**DOI:** 10.1101/635383

**Authors:** Matthew G. Marzo, Jacqueline M. Griswold, Kristina M. Ruff, Rachel E. Buchmeier, Colby P. Fees, Steven M. Markus

**Affiliations:** Department of Biochemistry and Molecular Biology, Colorado State University, Fort Collins, Colorado, USA; Department of Cell and Developmental Biology, University of Colorado School of Medicine, Aurora, United States

## Abstract

Cytoplasmic dynein plays critical roles within the developing and mature nervous systems, including effecting nuclear migration, and retrograde transport of various cargos. Unsurprisingly, mutations in dynein are causative of various developmental neuropathies and motor neuron diseases. These “dyneinopathies” define a broad spectrum of diseases with no known correlation between mutation identity and disease state. To overcome complications associated with studying dynein function in human cells, we employed budding yeast as a screening platform to characterize the motility properties of seventeen disease-correlated dynein mutants. Using this system, we have determined the molecular basis for several broad classes of etiologically related diseases. Moreover, by engineering compensatory mutations, we have alleviated the mutant phenotypes in two of these cases, one of which we confirmed with recombinant human dynein complexes. In addition to revealing molecular insight into dynein regulation, our data reveal an unexpected correlation between the degree of dynein dysfunction and disease type.

## INTRODUCTION

Motor-mediated intracellular transport is essential for numerous critical cellular processes^1–3^. This is especially apparent in motor neurons, in which cargoes must be transported over long distances to support various neuronal functions. For instance, the soma is the primary site of metabolic function where RNAs and proteins are synthesized and degraded. Thus, to maintain neuronal health, it is critical that cargoes are transported from the soma to the axon terminus (*i.e.*, along the axon) and vice versa. For example, neurofilaments, which provide structural stability to a cell, are transported to the axon terminus by plus end-directed microtubule motors (kinesins^4^) and to the soma by the minus end-directed microtubule motor, cytoplasmic dynein (hereafter referred to as dynein)^5–7^. Dynein has also been shown to play a key role in trafficking numerous other neuronal cargoes in various model organisms^5–18^, including autophagosomes^19–21^, mitochondria, and ionotropic glutamate receptors^22^. Defects or perturbations in dynein function lead to mislocalization of these cargoes, and results in an accumulation of protein aggregates at the neurite tip^19, 23^. In fact, accumulation of misfolded protein aggregates in the neuronal cytoplasm is a common pathological hallmark for motor neuron disease^5, 24, 25^. Consistent with an important role for dynein in neuronal health^26^, mouse models with dynein mutations exhibit severe neuropathy, and decreased rates of retrograde axonal transport, among other defects^27–29^.

In addition to its key role in the retrograde trafficking of cargoes, dynein also plays a critical and conserved role during neuronal development by promoting interkinetic neuronal migration (INM) in neuronal progenitor cells, and in the subsequent migration of young postmitotic neurons. During the former process, which is critical for neurogenesis, nuclear envelope anchored dynein motors promote migration of the nucleus from the basal to the apical surface of the neuroepithelium where mitotic divisions occur^30–32^. Thus, defects in dynein-mediated nuclear migration can lead to defects in early brain development.

Given its myriad roles in neuronal processes, it is unsurprising that mutations within the catalytic component of the dynein complex (the dynein heavy chain, or DHC, which is encoded by the DYNC1H1 gene) are found in individuals suffering from a wide array of neurodegenerative diseases. For instance, mutations in dynein underlie many cases of malformations of cortical development (MCD), spinal muscular atrophy with lower extremity dominance (SMA-LED), congenital muscular dystrophy (CMD), and Charcot-Marie-Tooth disease (CMT)^33–39^. Although mutations in the dynein regulator LIS1 are causative of the MCD disease lissencephaly^40–42^ (characterized by a smooth brain due to reduced or absent cortical folding), mutations in dynein more often lead to polymicrogyria, which is characterized by excessive small folds in the cerebral cortex^43–46^. Although a clear link has been established between dynein dysfunction and various neurological diseases, the underlying molecular basis for disease onset or progression is unknown.

An unambiguous mechanistic dissection of mutant dynein function in animal cells is complicated by the diverse cellular functions in which dynein participates (*e.g.*, axonal transport, centrosome separation, spindle assembly, nuclear envelope breakdown, spindle checkpoint inactivation)^47–52^, and the difficulties and expense associated with generating and analyzing mutant cell lines (*e.g.*, compromised viability, pleiotropism, heterozygosity). To overcome these issues, we have employed the versatility and power of the budding yeast *Saccharomyces cerevisiae* to understand how mutations found in individuals suffering from various neurological diseases lead to dynein dysfunction. In addition to their genetic amenability, low maintenance costs, and rapid generation time, the study of dynein function in budding yeast is simplified by several factors. In contrast to animal cells in which dynein performs numerous functions, the only known function for dynein in budding yeast is to position the mitotic spindle at the future site of cytokinesis^53–55^, making functional studies of dynein mutants in this organism simple and unambiguous. As in higher eukaryotes, the yeast dynein complex is comprised of light (Dyn2), light-intermediate (Dyn3), intermediate (Pac11), and heavy chains (Dyn1), the latter of which is the ATPase that powers motility along microtubules (see Fig. 1A)^56^. Whereas in humans, the non-catalytic subunits exist in different isoforms encoded by multiple genes and tissue-specific isoforms^57, 58^, each of the accessory chains in budding yeast is encoded by only a single gene, enabling simple genetic analysis and manipulation. Moreover, studies have revealed a high degree of structural similarity between yeast and human dynein^59, 60^, rendering structure-function studies in this organism relevant and translatable to animal cells. Compounded by the genetic amenability, ease of imaging, and the simple one-step method for isolation of recombinant, motile dynein motors^61–63^, budding yeast are a powerful model system for studies of dynein function.

**Figure 1.**
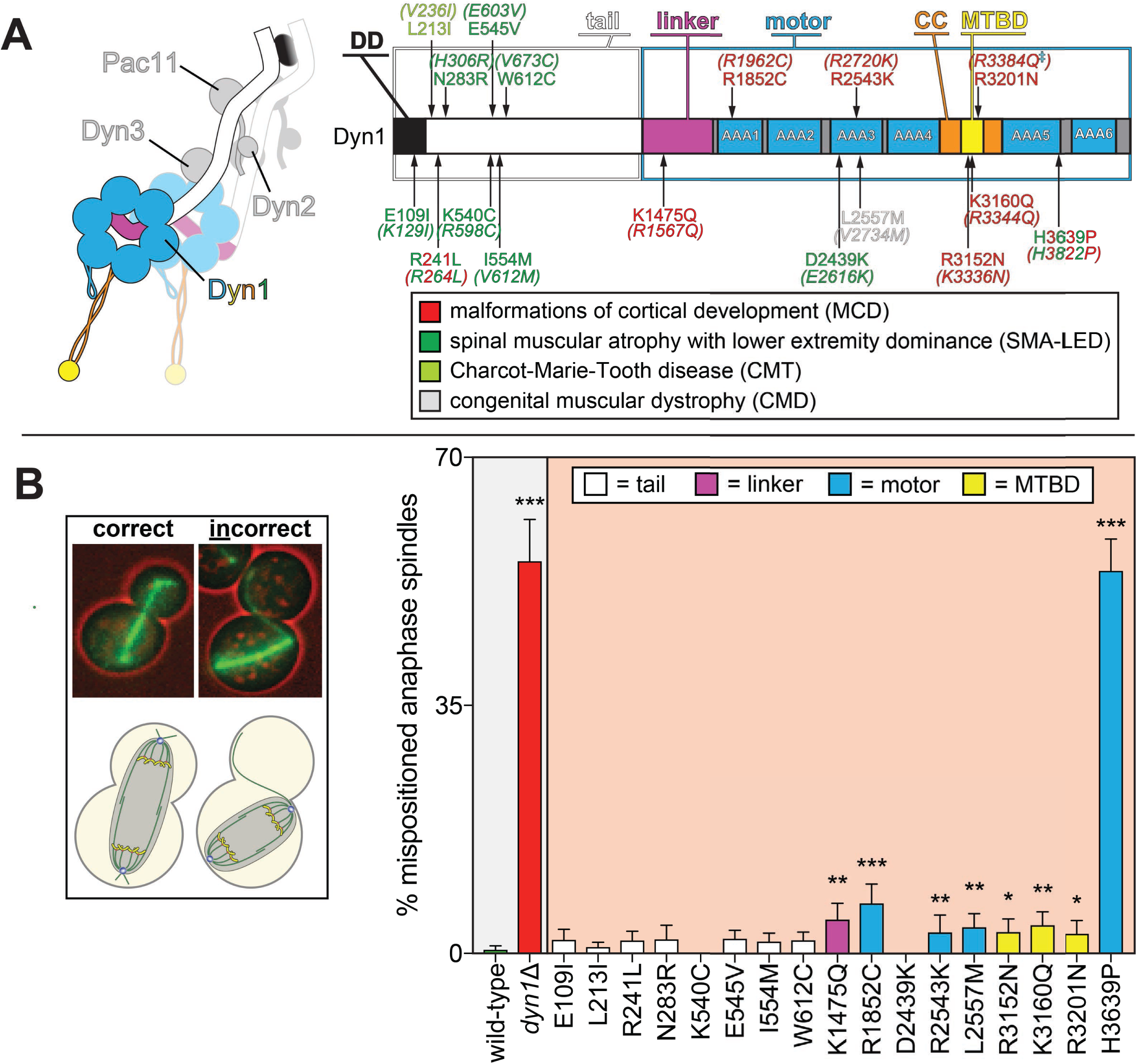
Spindle positioning assay provides coarse assessment of mutant dynein dysfunction. (A) Color-coded cartoon representation of the full-length dynein complex (left; with associated accessory chains; Dyn2, dynein light chain; Dyn3, dynein light-intermediate chain; Pac11, dynein intermediate chain; Dyn1, dynein heavy chain), and a linear schematic of Dyn1 with indicated disease-correlated mutations (right; DD, dimerization domain; CC, coiled-coil; MTBD, microtubule-binding domain). The equivalent human residues and disease-correlated substitutions are indicated in parentheses for each residue. ‡Note that we mistakenly substituted an asparagine for residue R3201 instead of a glutamine, the latter of which was identified as correlating with MCD^43^. R3201N was used throughout this study. (B) Representative fluorescence image (left; green, GFP-Tub1; red, contrast enhanced brightfield image to illustrate cell cortex) and quantitation of spindle positioning phenotypes in the 17 disease-correlated Dyn1 mutants, along with wild-type and dynein knock-out (*dyn1*Δ) cells. Each data point represents the fraction of mispositioned anaphase spindles along with standard error (weighted mean ± weighted standard error of proportion; n ≥ 99 anaphase spindles from three independent experiments for each strain). Statistical significance was determined by calculating Z scores, as described in the Materials and Methods (*, p ≤ 0.1; **, p ≤ 0.05; ***, p ≤ 0.001).

For this study, we focused on a library of 17 single point mutations in the dynein heavy chain (Dyn1), which are found in patients suffering from a broad spectrum of neurological diseases (Fig. 1A). These mutations were selected based on their conservation with corresponding residues in yeast dynein. Our findings reveal phenotypic signatures associated with the mutant library that range from partial to complete loss-of-function, and even gain-of-function in some respects. In some cases, the altered function was intrinsic to dynein, while in others, the effects could be attributed to alterations in the activity of the holo-dynein-dynactin complex. The rapid nature of our phenotypic analysis combined with the wealth of structural information available for dynein has enabled us to assess the likely structural basis for dysfunction in two instances. Moreover, our findings reveal an unexpected correlation between degree of dynein dysfunction and disease type. Overall, we describe a rapid and unambiguous set of tools that can be used to understand how disease-correlated mutations in dynein genes lead to onset or progression of disease.

## RESULTS

### Budding yeast as a model organism for dynein dysfunction

Genomic alterations in individuals suffering from a variety of neurological diseases have been mapped to numerous unique sites throughout the dynein heavy chain (Fig. 1A). The tail domain (~1400 amino acids) is the site of interaction for accessory chains (light-intermediate and intermediate chains), as well as adaptors that link dynein to the dynactin regulatory complex and various cellular cargoes^64^. The motor domain (~3000 amino acids) forms the catalytic core of dynein, where ATP binding and hydrolysis is translated into movement of the linker element that powers motility along microtubules^1, 59, 65^. Although the genetic basis for dyneinopathies is known, there exists very limited data on how these mutations are causative of motor dysfunction. Indeed, the mutations map throughout the entire heavy chain, with no clear correlation between disease state (*i.e.*, symptoms, severity, age of onset) and mutation identity. To understand how disease-correlated point mutations affect dynein function, we employed a series of well-established methodologies to assess the function of 17 single point mutants in budding yeast.

The first such assessment was to determine if the mutant motors were capable of correctly positioning the mitotic spindle, the only known function for dynein in budding yeast. To this end, we performed a spindle positioning assay using haploid yeast cells expressing GFP-Tub1 (α-tubulin; to visualize the mitotic spindle) and the mutant dynein from the native dynein locus. Given the dispensable nature of dynein function for yeast cell viability^53, 54^, a complete loss-of-function mutant would not be expected to compromise viability, but to simply affect spindle position. In this assay, single time-point images of mutant cells are acquired, and the position of the mitotic spindle is deemed to be either correct (*i.e.*, the anaphase spindle extended through the bud neck along the mother-bud axis) or incorrect (see Fig. 1B, left).

This analysis revealed that eight mutants exhibited varying degrees of spindle positioning defects that significantly differed from wild-type (Fig. 1B, right). All of those mutants with defects were those with substitutions in the motor domain, which encompasses the six AAA (<u>A</u>TPase <u>a</u>ssociated with various cellular <u>a</u>ctivities) domains, the linker element that performs the powerstroke, and the microtubule-binding domain (MTBD). All but one of the motor domain mutations led to significant defects. Seven out of the nine mutations linked to malformations in cortical development (MCD) were among those with defects in this assay, whereas none of the mutations associated with the other neurological diseases exhibited significant defects (see Discussion). Although most of these mutants differed to a small but significant degree from wild-type, H3639P exhibited defects as severe as loss of *DYN1* (*dyn1*Δ).

### Spindle tracking in live cells provides a sensitive read out for dynein-dynactin dysfunction

Given the somewhat binary nature of the spindle positioning assay, it provides only a coarse assessment of mutant functionality. Thus, as a more sensitive readout of mutant dynein function, we imaged dynein-mediated spindle movements in yeast cells and quantitatively assessed various parameters of these movements. Given the reliance of dynein on dynactin for this activity in cells, assessment of spindle movements is in fact a read-out of dynein-dynactin activity. To ensure that spindle translocation events were a consequence of dynein-dynactin activity, we performed these assays in cells deleted for *KAR9*, a genetic component of a pathway that promotes orientation of the spindle along the mother-bud axis^66–68^ (see Figure 2, Supplement 1). Moreover, we treated these cells with hydroxyurea (HU), an inhibitor of DNA synthesis that arrests yeast in a prometaphase-like state that precludes anaphase onset. This relatively simple but sensitive assay permits detection of subtle defects (or enhancements) in the motility parameters of dynein-dynactin^69^. In addition to obtaining velocity and displacement values, this assay also provides a read-out for relative activity (*i.e.*, how active dynein-dynactin is within the cell). Moreover, by scoring for “neck transit” success frequency – events in which a dynein-dynactin-mediated spindle translocation results in successfully transiting the narrow mother-bud neck – we are also able to determine if there are potential defects in force production. Previous studies have shown that neck transits are compromised in cases where dynein’s microtubule-binding affinity is weakened^70^, or when the CAP-gly domain of Nip100 (homolog of human p150 component of the dynactin complex) is genetically ablated^69^. Subsequent to HU arrest, full Z-stacks of the mitotic spindle and astral microtubules (via GFP-Tub1) were acquired every 10 seconds (see Fig. 2A for example), and the position of the spindle was subsequently tracked using a combination of manual (*e.g.*, to assess neck transit success) and automated 3-dimensional tracking (using a custom written Matlab-based routine).

**Figure 2.**
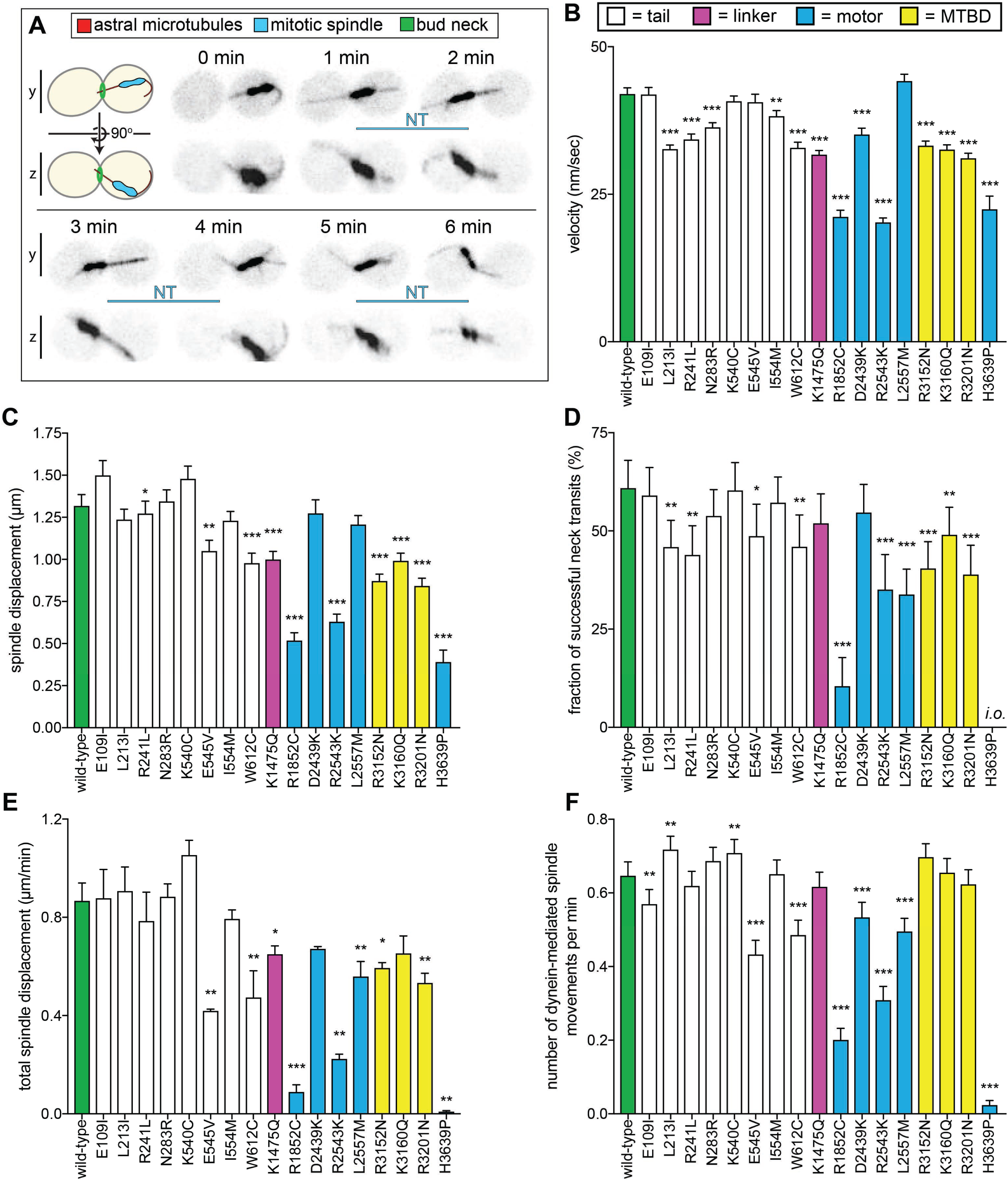
Quantitative assessment of dynein-dynactin-mediated spindle dynamics reveals refined insight into mutant dysfunction. (A) Cartoon and representative time-lapse fluorescence images of a hydroxyurea (HU)-arrested *kar9*Δ cell exhibiting typical dynein-mediated spindle movements, analysis of which is presented in panels B-F. Maximum intensity (X-Y projection; top) and Y-Z projections (bottom) are shown for each time point (NT, neck transit; note, line spans time frames over which the NT occurs). (B - F) Plots of indicated parameters for spindle dynamics in haploid wild-type and mutant strains. Briefly, the mitotic spindles were tracked in 3-dimensions using a custom written Matlab code. Dynein-mediated spindle movements were manually selected from the tracking data, from which velocity (B), displacement (C, per event; or, E, per minute), and the number of dynein-mediated spindle movements per minute (F) were obtained. The fraction of successful neck transits (successful attempts divided by total attempts) were manually scored (*i.o.*, insufficient observations; for H3639P, only 1 unsuccessful neck transit attempt was observed). Each data point represents the weighted mean ± weighted standard error (or standard error of proportion for D; n = 42 to 60 HU-arrested cells from three independent experiments were analyzed for each strain). Statistical significance was determined using an unpaired Welch’s t test (B and E), a Mann-Whitney test (C), or by calculating Z scores (D and F; *, p ≤ 0.1; **, p ≤ 0.05; ***, p ≤ 0.005). Also see Figure 2, Supplements 1, 2 3 and 4.

This analysis revealed a much more nuanced and broad array of defects in our library of mutants (Fig. 2B-F, and Fig. 2, Supplement 2A and B). Strikingly, all mutants exhibited varying degrees of alterations in their motility parameters with respect to wild-type cells. At the most severe end of the spectrum, H3639P cells – which also had the most severe spindle positioning phenotype – exhibited only 10 dynein-mediated spindle displacement events from all cells observed (compare to a mean of 244 events for all other strains; see Figure 2, Supplement 2A and B for scatter plots illustrating density of datasets). Consequently, this mutant had extremely low “activity” parameters (*i.e.*, total spindle displacement, Fig. 2E; and, number of dynein-mediated spindle movements per minute, Fig. 2F). The mutant with the second most severe spindle positioning phenotype, R1852C, exhibited significant defects in all motility metrics, including velocity, displacement (per event), neck transit success, and the two activity parameters. The relative phenotypic severity of these two mutants is consistent with findings from another group in which disease-correlated dynein mutants were assessed using recombinant human dynein complexes^71^. This group identified the human equivalents of R1852C (R1962C) and H3639P (H3822P) as being the most severe loss-of-function mutants in their reconstituted motility assays (see Discussion).

Another noteworthy mutant was K1475Q, a substitution within the linker domain, the mechanical element that is responsible for the powerstroke (Figure 2, Supplement 3). This mutation led to a reduction in spindle velocity, displacement, and also reduced the activity metrics of the motor. Given the position of this mutation, it may affect dynein activity by compromising linker remodeling during the priming or powerstroke movement of the linker. However, a recent study identified the equivalent human residue (R1567) as being at least partly required for the formation of an auto-inhibited conformation of human dynein called the phi-particle^72^ (due to its resemblance to the Greek letter^73^), thus raising the possibility that yeast dynein also adopts this conformation. Thus, the altered motility of this mutant may be a consequence of altered activity regulation (see Discussion).

All three mutations within the AAA3 module of the motor domain led to varying degrees of spindle motility alterations (D2439K, R2543K, and L2557M; Figure 2, Supplement 3). Interestingly, the most striking defect we observed with L2557M was a reduction in the neck transit success rate (Fig. 2D), which is the likely cause of the spindle positioning defect. This suggests that this mutation – which is buried within the large subdomain of AAA3, and makes hydrophobic contacts with a closely apposed helix – is likely affecting force generation by the motor.

All three microtubule-binding domain (MTBD) mutants exhibited fairly similar degrees and types of defects in effecting spindle movements (reductions in velocity, displacement, and neck transit success). Structural analysis revealed that all three mutations mapped to the surface of the MTBD that makes contacts with the microtubule (Figure 2, Supplement 3). Given all three substitutions lead to loss of a positive charge, it is likely that these mutations each lead to a reduction in the affinity of the motor for the negatively charged surface of the microtubule. The reduced activity metrics for R3152N and R3201N could thus be a reflection of a reduction in association kinetics of the mutants for the microtubule. This is supported by a study that found reduced microtubule binding for similar amino acid substitutions in a mouse dynein MTBD fragment^43^.

Although none of them led to significant spindle positioning defects, all of the tail domain mutations led to altered spindle motility parameters. Structural analysis revealed that most of these mutations (7 out of the 8) clustered to two distinct regions: (1) adjacent to, or within the N-terminal dimerization domain, or (2) at a surface that interfaces with the intermediate chain of a neighboring heavy chain in a 2 dynein:1 dynactin complex (Figure 2, Supplement 4). This latter region was recently identified as being important to stabilize the binding of a second dynein complex to dynactin, and ensuring that all four heavy chains are properly aligned for efficient motility of the human dynein-dynactin complex^74^. Our data suggest that the ability to recruit 2 dynein complexes to dynactin is conserved in yeast, and that disrupting this complex can compromise force generation (Fig. 2D) or activity (Fig. 2E and F). Although the last tail domain mutation, W612C, is in close proximity to this latter region, it is sufficiently far from the contact point with the second dynein complex to suggest that it is likely affecting some other facet of dynein-dynactin function.

### Mutations exert dominant negative effects on spindle dynamics

Given the heterozygous nature of these mutations in affected patients, we wondered how the mutants would behave in the presence of a second, wild-type copy of dynein. It is currently unclear whether cells with two copies of *DYN1* (*e.g.*, wild-type and mutant) assemble dynein complexes comprised of two different copies (*e.g.*, wild-type/mutant heterodimers), or whether they are comprised of only one copy (*e.g.*, wild-type homodimers, or mutant homodimers). This latter phenomenon could be a consequence of co-translational dimerization^75^. To distinguish between these two possibilities, we generated diploid yeast strains that contained one copy of a GFP-tagged dynein heavy chain (*DYN1-GFP*), and a second copy that was fused to an N-terminal affinity tag and a C-terminal HALO tag (*ZZ-DYN1-HALO*), the latter of which could be used to fluorescently label the motor (Fig. 3A). Lysate from these cells was subjected to affinity chromatography, and subsequent to incubation with a red fluorescent HALO ligand (HALO-TMR), the bound protein was eluted and used in a single molecule imaging experiment. If heterodimers assemble within cells, we expected to observe dual-color labeled molecules (green and red); however, if only homodimers form, then we expected to observe only red molecules (see Fig. 3A). Although we observed a small number of motile green molecules (0.9% of the total; likely due to contaminating Dyn1-GFP molecules in the protein preparation), and a single dual labeled molecule (0.3% of the total; Fig. 3B, arrow), the vast majority of motile molecules (98.8%) were exclusively red, indicating that dynein very rarely, if ever, forms heterodimers (Fig. 3C). This suggests a co-translational dimerization model for dynein complex assembly, similar to what has been observed for p53^76^.

**Figure 3.**
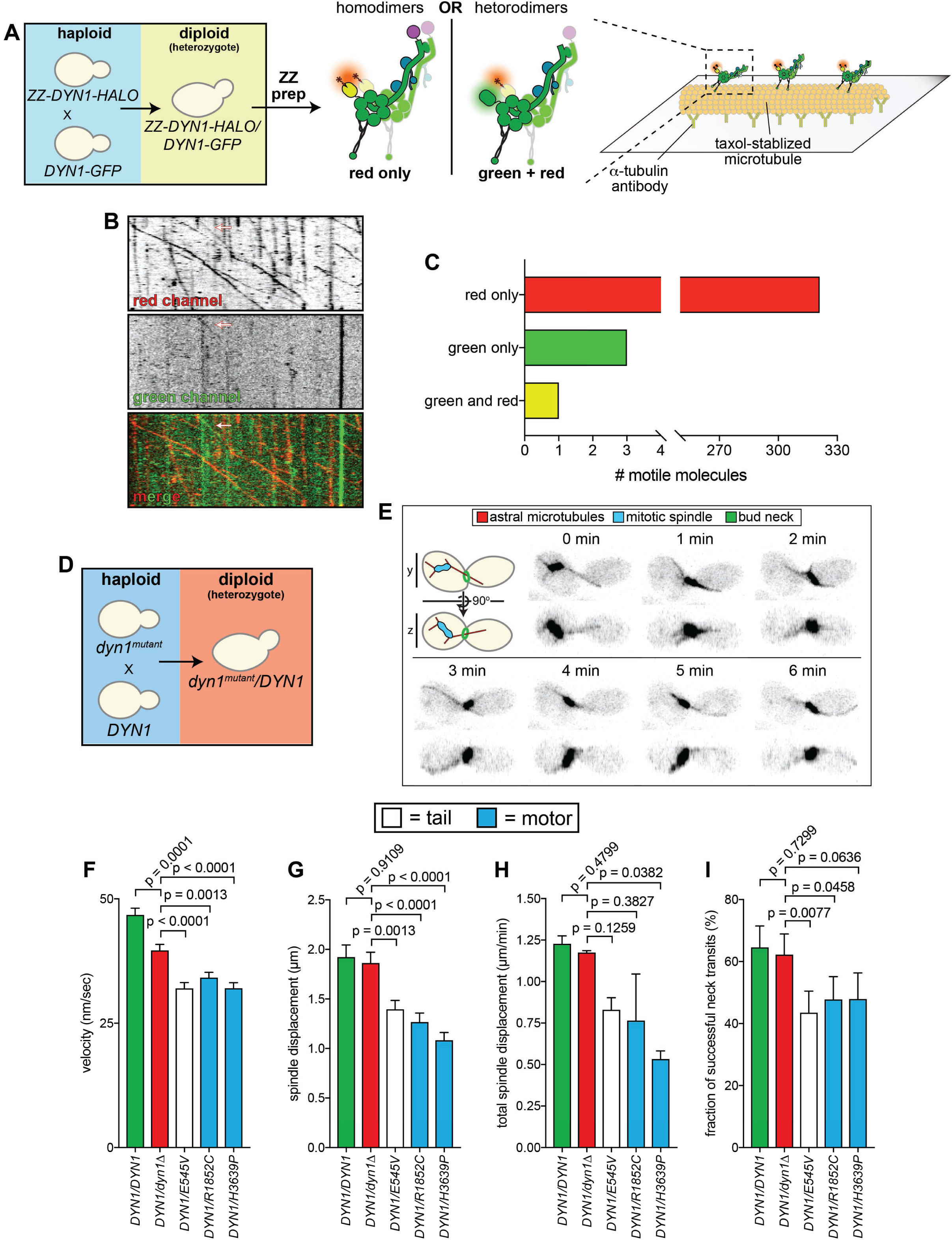
Quantitative assessment of spindle dynamics in heterozygous diploid cells reveals dominant nature of mutations. (A) Schematic depicting experimental approach to determine whether distinct protein from two different dynein alleles homo- or heterodimerize. (B and C) Representative kymograph (B) depicting large proportion of red (HALO-tagged) dynein molecules walking along microtubules (only one of which colocalized with a GFP-tagged dynein; arrow), along with associated quantitation (C). (D – E) Schematic depicting experimental approach to assess dynein-dynactin activity in heterozygous diploid cells (D), and representative fluorescence images of a diploid hydroxyurea (HU)-arrested *kar9*Δ/*kar9*Δ cell exhibiting typical dynein-mediated spindle movements. Maximum intensity (X-Y projection; top) and Y-Z projections (bottom) are shown for each time point. (F – I) Plots of indicated parameters for spindle dynamics in indicated diploid yeast strains. Each data point represents the weighted mean ± weighted standard error (or standard error of proportion for I; n ≥ 29 HU-arrested cells from two independent experiments were analyzed for each strain). Statistical significance was determined using an unpaired Welch’s t test (F and H), a Mann-Whitney test (G), or by calculating Z scores (I). Also see Figure 3, Supplement 1.

Next, we wished to recapitulate the heterozygous nature of some mutations in our budding yeast system. To this end, we chose three mutants – E545V, R1852C, H3639P – and determined whether they could affect dynein activity in heterozygous diploid yeast strains. We mated haploid wild-type cells with haploid mutant cells to generate heterozygous diploid cells (*e.g.*, *DYN1*/*dyn1^R^*^1852^*^C^*; Fig. 3D). In addition to comparing dynein activity in these cells to that from homozygous wild-type cells (*i.e.*, *DYN1*/*DYN1*), we also compared them to hemizygous cells with only one copy of *DYN1* (*i.e.*, *DYN1*/*dyn1Δ*; Fig. 3E-I, and Fig. 3, Supplement 1). Although homozygous wild-type cells exhibited similar dynein-dynactin activity to the hemizygotes, the velocity was somewhat reduced in the latter, suggesting a critical concentration of dynein is required for effecting maximal spindle velocity (Fig. 3F, and Fig. 3, Supplement 1). Analysis of the heterozygous mutants revealed that all three exhibited partial loss-of-function phenotypes in almost all assays with respect to the hemizygous cells, indicating that they are all indeed dominant alleles. Taken together, these data indicate that homodimers of mutant dynein are sufficient to compromise the activity of wild-type homodimers within the cell.

### Single molecule motility assays reveal insight into dynein-intrinsic dysfunction

Findings from our spindle tracking assay revealed the consequences of mutations on various parameters of dynein-dynactin-mediated spindle movements. These movements are mediated by cortically anchored dynein-dynactin complexes that are regulated at various levels by numerous effector molecules (*e.g.*, Pac1, Ndl1, Num1, She1^62, 77–79^). For instance, cortical targeting of dynein is affected by various molecules, including Pac1, which tethers dynein to microtubule plus ends^80^, and Num1, which anchors dynein-dynactin complexes to the cortex^81^. Thus, mutations that alter the ability of dynein to interact with or be affected by these molecules will result in alterations in dynein-dynactin-mediated spindle movements. To determine whether mutations affect dynein-intrinsic activities, it is therefore important to specifically assess dynein activity without these complicating factors. To this end, we employed a single molecule motility assay, in which the movement of individual purified dynein motors is quantitatively assessed. Performing this assay using yeast dynein has one key advantage over human dynein: unlike human dynein, which requires dynactin and one of several adaptor molecules for processive single molecule motility (*e.g.*, BicD2, Hook3^82, 83^), yeast dynein is a processive motor without these factors^61^. This somewhat unique property of yeast dynein thus permits an unbiased assessment of dynein-intrinsic motility.

Single molecule motility analysis of each mutant revealed that thirteen of those that exhibited defects in the spindle tracking assay showed some degree of motility alteration in the *in vitro* assay (Fig. 4; see Figure 4, Supplement 1 for scatter plots and some example kymographs). Interestingly, the precise defect *in vivo* was not always predictive of the alteration in dynein motility *in vitro*. Although the reasons for this are unclear, they are likely due in part to the differences in requirements for spindle transport versus those for unloaded single molecule motility (*i.e.*, in which there is no resistance to transport). Additionally, small defects in single motor motility may lead to more pronounced defects in the context of a motor ensemble, as may be the case for dynein at the cell cortex^77^. Such mutants included W612C and R1852C, both of which led to a reduction in all spindle tracking metrics in cells, but had very little effect on the displacement (run length) of single molecules *in vitro*. Similarly, R2543K, which reduced spindle velocity by ~50% had only a minor effect on single molecule velocity values.

**Figure 4.**
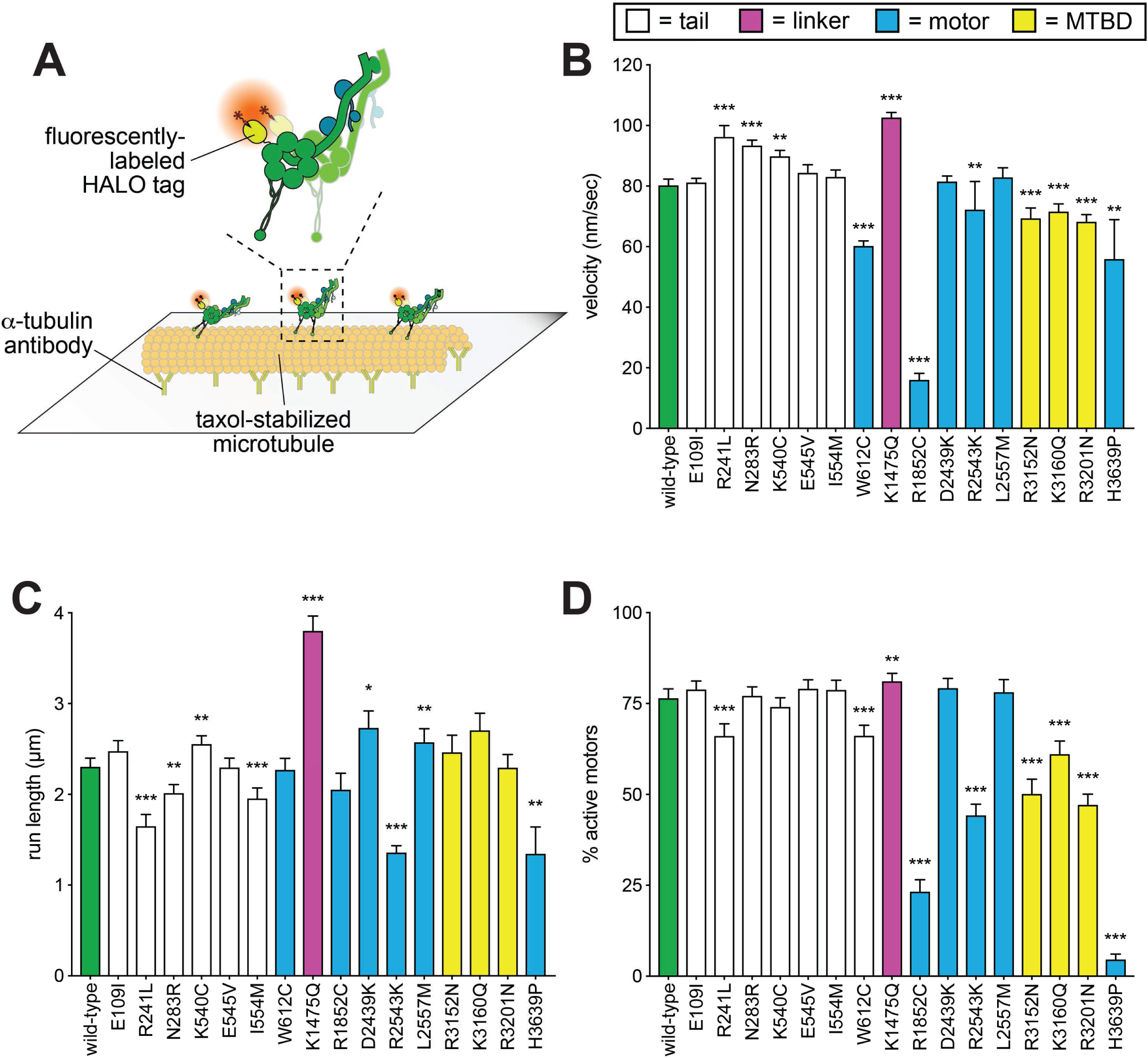
Single molecule analysis reveals insight into dynein-intrinsic dysfunction. (A) Cartoon representation of experimental approach. (B – D) Quantitation of indicated parameters of single molecule motility. Each data point represents the weighted mean ± weighted standard error (or standard error of proportion for D; n ≥ 284 single molecules from at least two independent experiments were analyzed for each motor variant). Technical difficulties precluded us from generating the L213I mutant in the protein purification background. Statistical significance was determined using an unpaired Welch’s t test (B), a Mann-Whitney test (C) or by calculating Z scores (D; *, p ≤ 0.1; **, p ≤ 0.05; ***, p ≤ 0.005). Also see Figure 4, Supplement 1.

In a few cases, we observed little or no difference from wild-type in single molecule motility parameters in spite of differences in the spindle tracking assay (*i.e.*, E109I, K540C, E545V, I554M, D2439K, L2557M). In two of these cases (E545V and L2557M), one of these changes was a reduction in the neck transit success rate (see Fig. 2D). As discussed above, this phenomenon is potentially a read-out of force generation. Given the unloaded nature of single molecule motility, this assay would be unable to detect differences in force generation by dynein. Although the molecular determinants of dynein force production are not well understood, it is possible that these mutations specifically affect the ability of dynein to remain bound to microtubules under conditions of high load.

Although most mutants exhibited loss-of-function phenotypes in the *in vitro* assay, a few mutations led to gain-of-functions. For instance, R241L, N283R, and K1475Q all led to an increase in velocity, while K1475Q also caused an increase in run length and the fraction of active motors (see Figure 4, Supplement 1C for example kymograph). We confirmed the increased run length for K1475Q was not a consequence of motor aggregates, which could presumably lead to increased processivity^84^ (Figure 4, Supplement 2). As mentioned above, R241 and N283 are adjacent to the N-terminal dimerization domain, and K1475 is within the linker domain (see Figure 2, Supplements 3 and 4). In the cases of R241L and N283R, the gain-of-functions observed *in vitro* are possibly causative of the *in vivo* deficiencies, indicating that a faster or more processive motor is not advantageous for the spindle positioning function (see Discussion regarding K1475Q). This is consistent with recent studies that showed gain-of-functions in dynein (or dynein regulators) can lead to defects in dynein-mediated neuronal maturation^85^, or spindle orientation^72^.

In summary, in all cases in which motility parameters were altered *in vitro* as a consequence of a particular mutation, we are able to conclude that the underlying molecular defect is most likely intrinsic to dynein itself, and is likely not a consequence of an altered interaction with either accessories or regulators.

### Localization phenotypes provide mechanistic basis for dysfunction in a subset of mutants

Our *in vivo* and *in vitro* functional data described above revealed the particular parameters of dynein motility that were altered by the mutations, and also whether the altered motility was in fact intrinsic to dynein (*i.e.*, if there was an *in vitro* phenotype), or was possibly a consequence of alterations in interactions with regulators such as dynactin. As discussed above, proper dynein function in yeast relies on the coordinated effort of various molecules to localize dynein to microtubule plus ends, from where it is offloaded to its site of action: the cell cortex (Fig. 5A)^63^. For instance, dynein plus end localization occurs in a dynactin-independent manner^86^, but requires an interaction with Pac1 (the LIS1 homolog)^80^, as well as the accessory chains Dyn3^56^ (light-intermediate chain) and Pac11 (intermediate chain)^87^. In contrast, dynactin is required for dynein to localize to cortical Num1 receptor sites^86^. Thus, quantitative assessment of dynein localization can reveal potentially altered interactions with various regulators or accessories.

**Figure 5.**
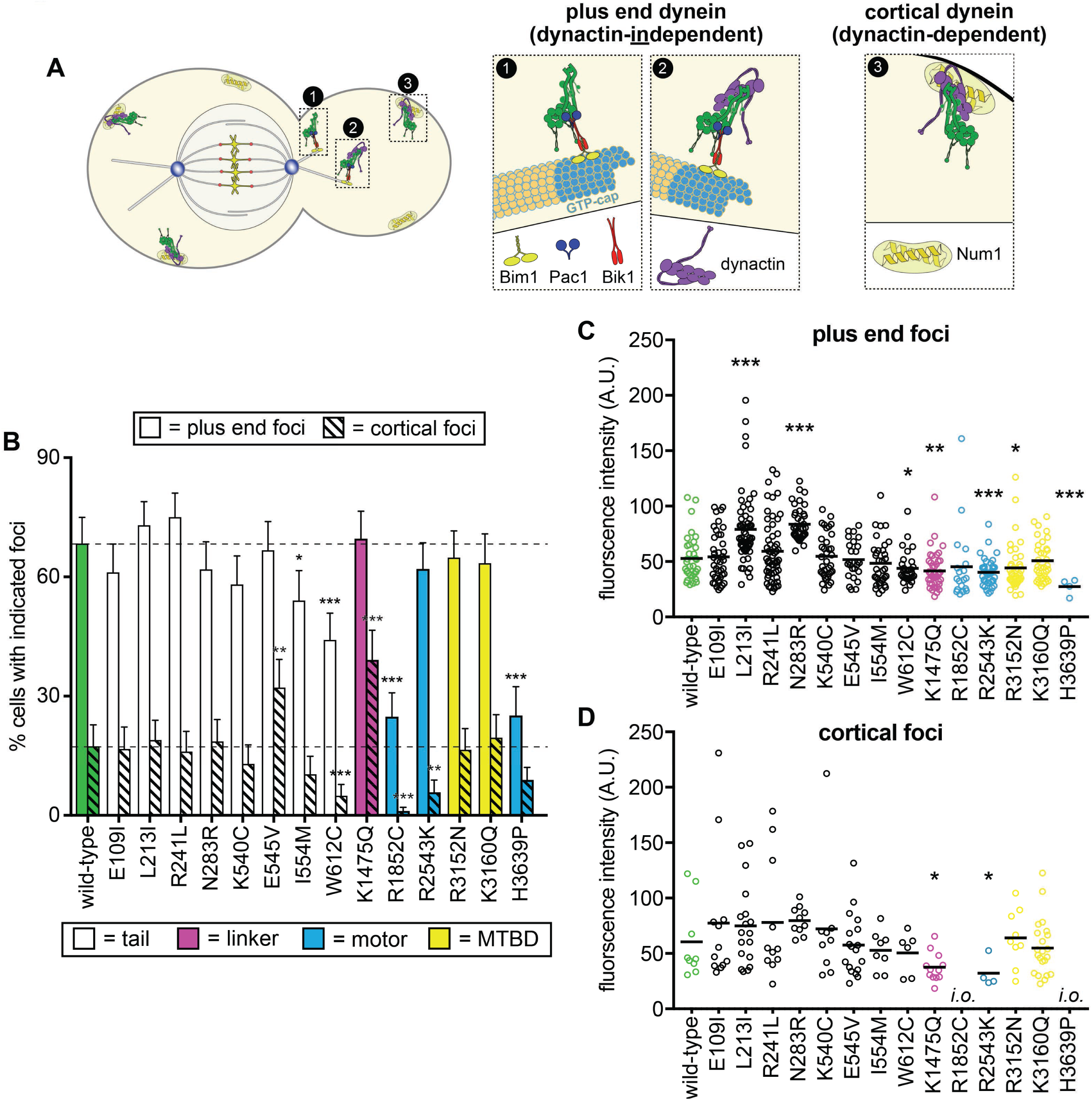
Quantitative assessment of dynein localization reveals potential basis for mutant dysfunction. (A) Cartoon representation depicting the two main sites of dynein localization, and the molecular basis for each. Dynein plus end localization (1) requires Bik1, Pac1, and possibly Bim1, but does not require dynactin. Rather, dynactin plus end localization (2) relies on dynein. Association of dynein with the cortex requires (3) dynactin and the cortical receptor, Num1. (B) The frequency of dynein localization to either microtubule plus ends or the cell cortex is plotted for indicated strains. To enrich for mitotic cells, overnight cultures were diluted into fresh media for 1.5 hours prior to imaging. To further reduce variability due to cell cycle-dependent changes^88^, localization frequency was scored for mitotic cells only. Each data point represents the weighted mean ± weighted standard error (68 to 111 mitotic cells from at least two independent experiments were analyzed for each strain). (C and D) Fluorescence intensity values for either plus end (C) or cortical (D) dynein foci observed in mitotic cells described in B (4 to 59 plus end foci, and 4 to 22 cortical foci from two independent experiments were analyzed; *i.o.*, insufficient observations; only 1 cortical focus was observed for both R1852C and H3639P). Statistical significance was determined by calculating Z scores (B), or by applying an unpaired Welch’s t test (C and D; *, p ≤ 0.1; **, p ≤ 0.05; ***, p ≤ 0.005).

For this analysis, we focused on a select group of mutants: those with substitutions within the N-terminal tail domain (which is the site for interaction with the accessory chains, dynactin, and Num1^88^), those with the most severe phenotypes (R1852C, R2543K, and H3639P), and two of the MTBD mutants. We acquired time-lapse images of haploid cells expressing fluorescently-labeled tubulin (mRuby2-Tub1; α-tubulin) and 3GFP-tagged copies of each dynein mutant. Plus end and cortical foci were identified from movies, and their targeting frequency (*i.e.*, percent cells with foci) and fluorescence intensities (a readout of molecule number per site) were quantified.

This analysis revealed significantly altered localization frequencies or intensities for most of the mutants, including L213I, N283R, E545V, I554M, W612C, K1475Q, R1852C, R2543K, and H3639P (Fig. 5A-D). For instance, the K1475Q substitution (within the linker) led to a large increase in the frequency of observing cortical foci (2.2-fold; p = 0.0011), but a concomitant reduction in the number of molecules per cortical focus (38% reduction in fluorescence intensity; p = 0.0117), whereas W612C, R1852C and H3639P all exhibited a reduction in dynein levels at plus end and cortical sites (see Discussion).

### Proline-dependent structural constraint is the likely cause for dysfunction in dynein H3639P

Given the wealth of structural information currently available for dynein, we sought to determine the structural basis for dysfunction in two of the mutants: R1852C and H3639P. Analysis of available dynein structures revealed that H3639 is situated within a conserved loop that connects two alpha helices, one of which is an extension of the buttress, a structural element that helps to communicate nucleotide-dependent structural rearrangements within the AAA ring to the MTBD (Fig. 6A)^65^. We first asked whether gain of proline or loss of histidine is the cause for the severe loss-of-function. To this end, we substituted H3639 with either a serine (to preserve the polar nature and hydrogen-bonding capabilities of histidine), valine (a hydrophobic residue), or asparagine (found in the equivalent position of human dynein-2), and assessed the activity of these mutants in the spindle positioning assay. None of these substitutions led to significant spindle positioning defects, indicating that gain of proline is the reason for dynein dysfunction in H3639P (Fig. 6B). We next asked if proline substitutions are tolerated at other sites within this inter-helical loop. Using the spindle positioning assay, we found that proline was well tolerated at all sites within the inter-helical loop except for position 3641 (2 residues C-terminal to 3639; Fig. 6B). Thus, introduction of a proline at two distinct sites within this loop are sufficient to severely compromise dynein function.

**Figure 6.**
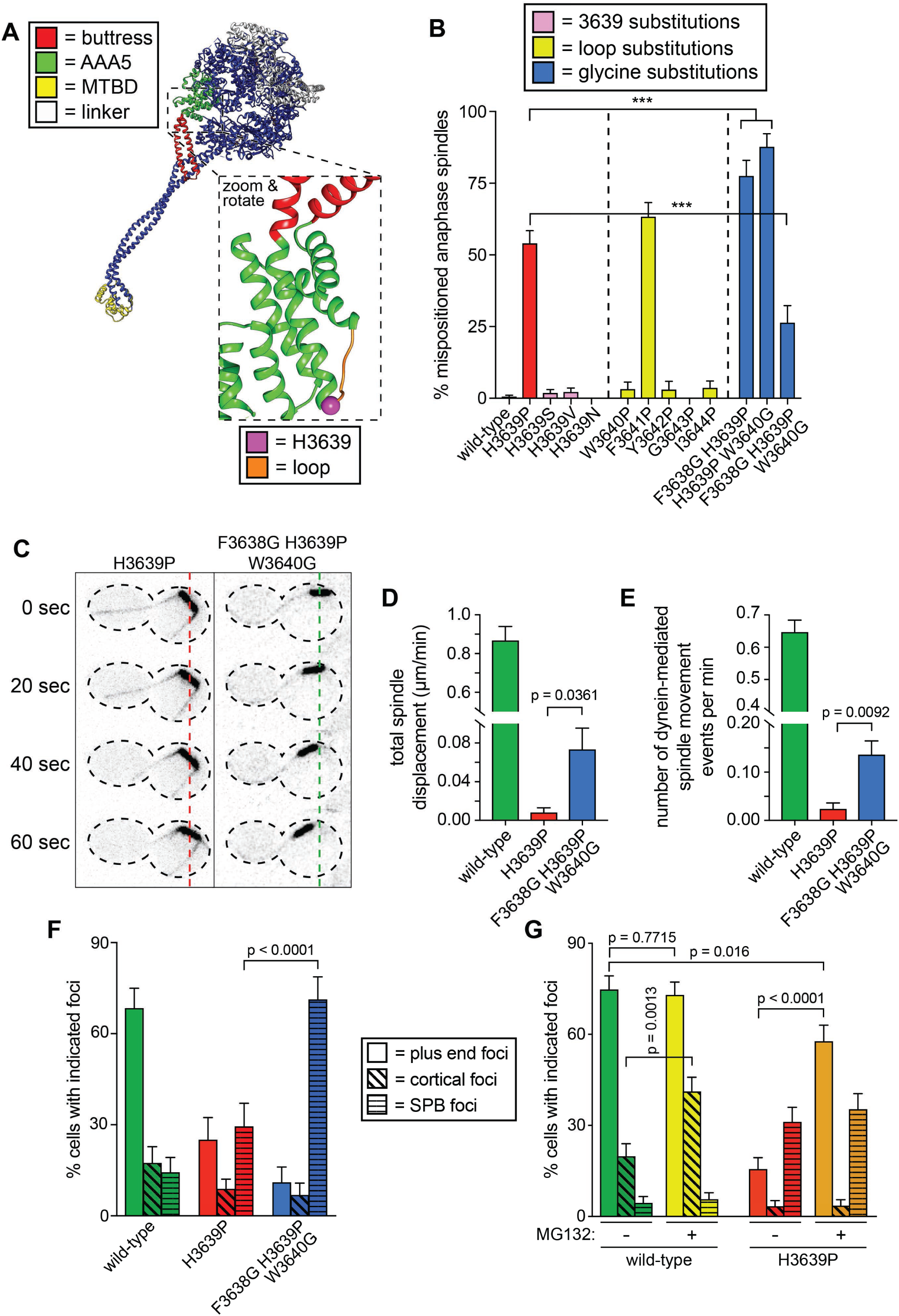
Detailed dissection of the molecular basis for dysfunction in H3639P. (A) Color-coded structural model of the dynein motor domain (from PDB 4RH7^65^) with zoomed in region depicting H3639 residing within an inter-helical loop within AAA5. (B) Fraction of cells with mispositioned spindles are plotted for yeast strains with indicated dynein mutations. Each data point represents the fraction of mispositioned anaphase spindles along with standard error (weighted mean ± weighted standard error of proportion; n ≥ 67 anaphase spindles from at least two independent experiments for each strain; *, p ≤ 0.1; **, p ≤ 0.05; ***, p ≤ 0.005). (C) Representative time-lapse fluorescence images of two indicated hydroxyurea (HU)-arrested *kar9*Δ cells. Note the lack of spindle translocation in H3639P, but the clear dynein-mediated movement in the F3638G H3639P W3640G (dashed lines provide a point of reference). Maximum intensity projections are shown for each time point. (D - E) Plots of two activity parameters for spindle dynamics in indicated haploid strains (total displacement, D; and, number of events per minute, E). Each data point represents the weighted mean ± weighted standard error (D) or standard error of proportion (E; n ≥ 29 HU-arrested cells from at least two independent experiments were analyzed for each strain). (F and G) The frequency of dynein localization to either microtubule plus ends, the cell cortex, or spindle pole bodies (SPBs) is plotted for indicated strains and drug treatment (for mitotic cells only). In addition to the indicated alleles and drug treatment, the plot in panel G depicts cells that possess the *prd1-DBD-CYC8* allele, which represses transcription of pleiotropic drug resistance genes^91^, thus promoting intracellular retention of MG132. These cells were treated with 75 µM MG132 for 1.5 hours prior to imaging (control cells were treated with an equal volume of DMSO). Each data point represents the weighted mean ± weighted standard error (73 to 107 mitotic cells from two independent experiments were analyzed for each strain). Statistical significance was determined by calculating Z scores (B, E, F and G), or by applying an unpaired t-test with Welch’s correction (D). Also see Figure 6, Supplement 1.

We hypothesized that prolines are not tolerated in these two regions (residues 3639 and 3641) of the inter-helical loop because of the structural constraints imposed by prolines due to bond angle restrictions. If this were true, then we reasoned that increasing the structural flexibility in the immediate vicinity of P3639 might rescue the proline-dependent defects. To test this, we introduced glycine substitutions at either the N-terminal residue (F3638), the C-terminal residue (W3640), or both. For reasons that are unclear, both double mutants (*i.e.*, F3638G H3639P, and H3639P W3640G) led to spindle positioning defects that were significantly more severe than H3639P and dynein knock-out cells (*dyn1Δ*; p < 0.0001; Fig. 6B). Strikingly, however, the triple mutant – in which P3639 is flanked by glycines (*i.e.*, F3638G H3639P W3640G) – exhibited spindle positioning defects that were significantly less severe than H3639P (p < 0.0001).

To confirm these findings, we assessed the activity of the triple mutant using the spindle tracking assay. Although the activity of this mutant was much less than that of wild-type dynein-expressing cells, it was significantly greater than the H3639P single mutant (p ≤ 0.0361; Fig. 6C-E). In spite of the increase in activity, the quality of the spindle movements (velocity and displacement per event) were nearly identical between H3639P and the triple mutant (Figure 6, Supplement 1A - D). Interestingly, the single molecule assay revealed that the triple mutant was no better than the single mutant in any metrics (Figure 6, Supplement 1E - I). This indicates that the triple mutant does not rescue dynein motility, but some other metric of *in vivo* dynein (or dynein-dynactin) activity. We next imaged dynein in live cells to determine if the triple mutant alters any aspects of dynein localization. Although the flanking glycines did not rescue plus end or cortical localization, it did lead to a large increase in the fraction of cells exhibiting dynein foci at the spindle pole bodies (SPBs; Fig. 6F; p < 0.0001), suggesting that the triple mutant increases the relative concentration of dynein within cells with respect to the single H3639P. The relevance of dynein localization to the SPB is unclear, but previous work from our lab demonstrate that this localization requires the MTBD^79^. In light of our other observations, the increased localization frequency of the triple mutant suggests that the flanking glycines rescue H3639P cellular defects by reducing proline-dependent inflexibility, which in turn prevents a protein degradation response, thus increasing the relative concentration of dynein within the cell. To test whether protein degradation is indeed one of the underlying means for dysfunction of H3639P, we assessed dynein localization in the absence or presence of MG132, a proteasome inhibitor. Addition of MG132 increased the frequency of wild-type cortical dynein foci by approximately 2-fold (p = 0.0013), but had no significant effect on the frequency of either SPB or plus end targeting (Fig. 6G). Consistent with our hypothesis, MG132 treatment increased the frequency of plus end targeting of H3639P by 3.7-fold (p < 0.0001) to levels comparable to wild-type dynein; however, the frequency of cortical targeting of H3639P was unaffected by proteasome inhibition.

### An ectopic disulfide bond is the likely cause for R1852C dysfunction

Close inspection revealed a cysteine situated within very close proximity to R1852 (~3 Å; Fig. 7A), which lies within the first AAA module (AAA1). Given the mutation results in a cysteine substitution at this site, we reasoned that an ectopic disulfide bond may be responsible for the phenotypic consequences. To determine whether this was the case, we mutated the closely apposed, highly conserved cysteine to a serine (C1822S; Fig. 7B), which would eliminate the potential for disulfide bond formation at this site. We then used several of our assays to quantitate the degree to which C1822S might rescue defects due to R1852C.

**Figure 7.**
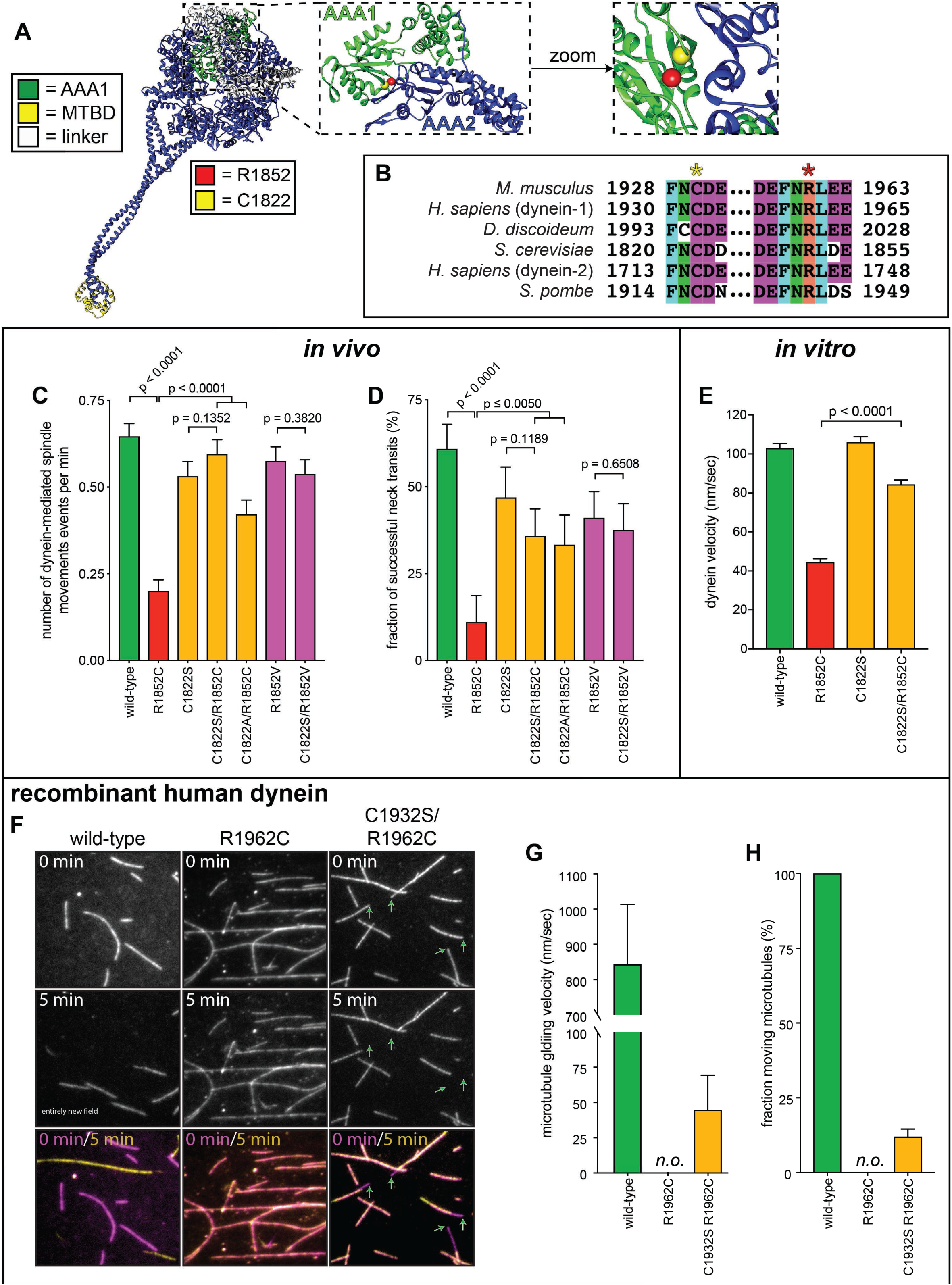
Detailed dissection of the molecular basis for dysfunction in R1852C. (A) Color-coded structural model of the dynein motor domain (from PDB 4RH7^65^) with zoomed in region depicting R1852 residing within AAA1 near its interface with AAA2. (B) Sequence alignment illustrating the high degree of conservation for C1822 (yellow star) and R1852 (red star) among various dynein heavy chains. (C and D) Plots of two activity parameters for spindle dynamics in indicated haploid strains (total displacement, C; and, fraction of successful neck transits, D). Each data point represents the weighted mean ± weighted standard error (C) or standard error of proportion (D; n ≥ 28 HU-arrested cells from two independent experiments were analyzed for each strain). (E) Plot of velocity values for single molecules of indicated GST-dynein_331_ variants (n ≥ 158 single molecules from at least two independent experiments were analyzed for each motor variant). (F) Representative fields of microtubules being translocated by surface-adsorbed recombinant human dynein complexes. (G and H) Plots depicting velocity (G) and fraction (H) of microtubules (at least 50 microtubules from two independent experiments were analyzed for each variant; “*n.o.*”, none observed). Statistical significance was determined by calculating Z scores (B and E), or by applying an unpaired Welch’s t test (D). Also see Figure 7, Supplement 1.

Although C1822S alone compromised all parameters of dynein-dynactin-mediated spindle movements, this substitution led to a partial restoration of most of the parameters in the R1852C mutant (*i.e.*, C1822S R1852C double mutant; Fig. 7C and D, and Figure 7, Supplement 1A-C). We observed a similar restoration of function in a C1822A R1852C double mutant. We ruled out the possibility that the partially hydrophobic nature of cysteine is the cause for the dysfunction in R1852C by assessing an R1852V mutant, which exhibited defects that were significantly less severe than R1852C. Moreover, combining C1822S with R1852V led to no degree of rescue with respect to the single R1852V mutant, which is in stark contrast to our observations with the double C1822S R1852C mutant (Fig. 7C and D, and Figure 7, Supplement 1A-C). Using our localization assay, we found that C1822S rescued the localization of the R1852C mutant to plus ends and the cell cortex almost to wild-type levels (Figure 7, Supplement 1D). As with the spindle tracking assay, although R1852V led to localization defects, the C1822S R1852V double mutant was almost identical to R1852V alone.

Before assessing the extent of rescue with the single molecule assay, we first tested whether using a minimal dynein fragment that is sufficient for processive motility^61^ would rescue any of the motility defects we observed with the R1852C mutant (GST-dynein_331_). This fragment – a glutathione S-transferase (GST)-dimerized motor domain that exhibits motility parameters that are very similar to the full-length molecule^61^ – lacks the tail domain, and thus does not co-purify with or rely on any of the accessory chains. If protein misfolding and/or degradation is partly to blame for mutant dysfunction – as suggested from the single molecule (only 23.2% active motors; Fig. 4D) and localization assays (severe reduction in localization frequency; Fig. 5B) – then we reasoned that the simplicity and compact fold of GST-dynein_331_ might rescue some of these defects. In striking contrast to the full-length mutant, the fraction of active GST-dynein_331_ R1852C motors was almost identical to that of the wild-type GST-dynein_331_ motor (note the same construct did not rescue the H3639P mutant; Figure 7, Supplement 1E). Although the reasons for this are unclear, they.. However, much like the full-length molecule, the velocity of the GST-dynein_331_ mutant was severely reduced with respect to wild-type (Fig. 7E and Figure 7, Supplement 1G). Consistent with the spindle tracking data, C1822S was indeed sufficient to significantly rescue the single molecule velocity defect in the minimal dynein fragment (Fig. 7E and Figure 7, Supplement 1G). Taken together, these findings indicate that an ectopic disulfide bond is the likely cause for R1852C dysfunction.

### Compensatory mutation partially suppresses motility defects of human dynein mutant

Finally, we sought to determine whether the compensatory cysteine to serine substitution would also rescue motility defects in human dynein. To this end, we engineered the equivalent mutations (C1932S, R1962C, or C1932S R1962C) into an insect cell expression plasmid encoding an affinity-tagged human dynein complex (ZZ-SNAPf-DYNC1H1, IC2, LIC2, Tctex1, Robl1, LC8). Subsequent to purification of recombinant complexes, we assessed their function using a microtubule gliding assay in which free microtubules are translocated by dynein complexes non-specifically adsorbed to the glass surface. This assay permits an assessment of dynein-intrinsic motility parameters without the need for additional factors that are required for single molecule motility of human dynein (*i.e.*, dynactin and a requisite adaptor^82, 83^). Consistent with previous findings, the single R1962C mutation (equivalent to R1852C) was sufficient to completely disrupt microtubule gliding activity^71^ (Fig. 7F-H); however, as with yeast dynein, substitution of the closely apposed cysteine to a serine (C1932S) was sufficient to restore some of the lost function. Although the compensatory mutation did not restore activity to wild-type levels (note the low velocity and degree of activity with respect to wild-type in Fig. 7G and H), the same was true for the full-length yeast dynein complex, which was only partially rescued by the mutation (see Figs. 7C and D, and Figure 7, Supplement 1A-C). Although unclear, the different degrees of rescue by the compensatory mutation with yeast dynein (observed *in vivo*; Fig. 7C and D, and Figure 7, Supplement 1A-D) versus human dynein (observed *in vitro*; Fig. F-H) may be due to the presence of quality control mechanisms at play in live yeast cells that ensure only properly folded motors are delivered to cortical receptor sites. No such mechanism is present to prevent adsorption of inactive recombinantly produced human dynein motors to the glass in the *in vitro* gliding assay. In summary, these results confirm our findings with yeast dynein, and moreover validates yeast as a powerful model system to understand the molecular basis for dynein dysfunction in patients.

### Severity of dynein dysfunction correlates with disease type

In an effort to identify potential correlations between degree of dynein dysfunction and disease, we summarized the findings from all of our various assessments into a heat map in which the variance (or lack thereof) from wild-type was assigned a color based on the statistical significance (*e.g.*, green, p ≥ 0.100; red, p ≤ 0.005; Fig. 8A). Unsurprisingly, on average, mutations within the highly conserved motor domain exhibited more statistically significant differences from wild-type than those in the tail domain. Moreover, this broad view of dysfunction in the mutant library revealed that the more severe loss-of-function mutants appeared to correlate with MCD, while those with lesser defects mainly correlate motor neuron diseases (SMA-LED, CMT or CMD).

**Figure 8.**
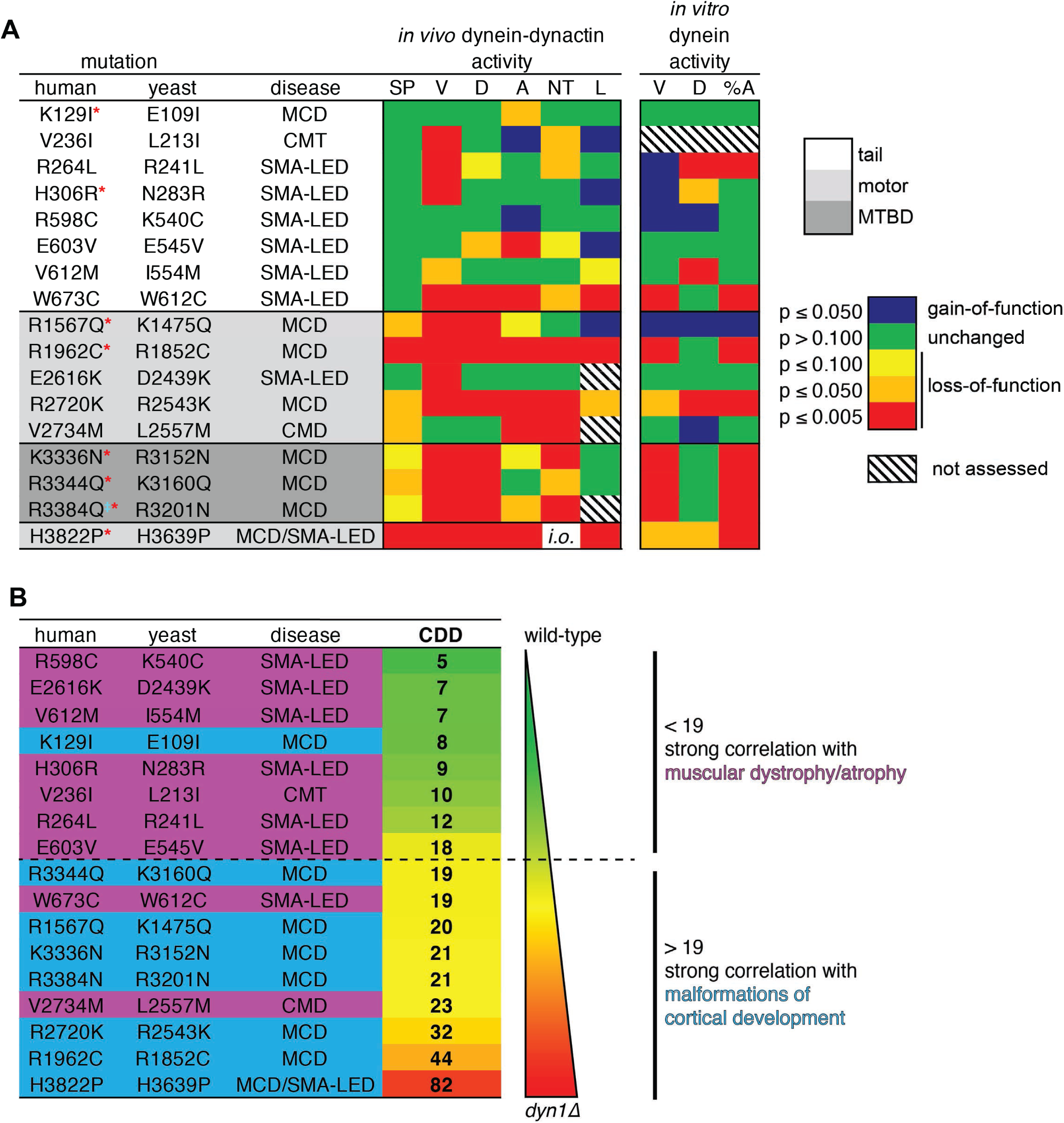
Summary of phenotypic analysis of disease-correlated mutants. (A) Heat map depicting degree of statistical significance for difference between each mutant and wild-type cells for the indicated assays (SP, spindle positioning; V, velocity; D, displacement per event; A, activity; NT, neck transit; L, localization; %A, percent active motors; MCD, malformations in cortical development; CMT, Charcot-Marie-Tooth disease; SMA-LED, spinal muscular atrophy with lower extremity dominance; CMD, congenital muscular dystrophy). Red asterisks depict mutants that were assessed in previous study using recombinant human dynein^71^. ‡Note that we mistakenly substituted an asparagine for residue R3201 instead of a glutamine, the latter of which was identified as correlating with MCD^43^. R3201N was used throughout this study. Deviation from wild-type cells in either the “total spindle displacement” (see Fig. 2E), or the “number of dynein-mediated spindle movements per minute” (see Fig. 2F) metric was used for the activity column (“A”). Mutants are listed from N- to C-terminus of Dyn1 (shading indicates in which domain of Dyn1 each mutation resides). Significance was calculated as indicated in previous figure legends. (B) Each mutant ranked by their coefficient of dynein dysfunction score (CDD; see text and Figure 8, Supplement 1). As shown, above a CDD score of ~18, the correlation with malformations in cortical development increases, while that with motor neuron diseases decreases. Also see Figure 8, Supplement 1.

To more quantitatively assess our data for potential correlations between dynein dysfunction and disease type, we developed a system in which the degree of variance from wild-type for each mutant were assigned numerical scores that we then used to tabulate a single value that represents the degree of dynein dysfunction for each mutation (coefficient of dynein dysfunction, or CDD). For tabulation of the CDD value, we focused on the *in vivo* data only. The reasons for this were two-fold: (1) defects observed in the *in vivo* assays were largely reflective of the degree of dysfunction, whereas the *in vitro* data generally revealed the mechanisms of dysfunction (*e.g.*, whether the mutation affected dynein-intrinsic or extrinsic function); and, (2) all mutants with *in vivo* defects also exhibited *in vitro* defects, and so we chose to include only the *in vivo* data to avoid redundancies in the CDD tabulation. In addition to the spindle positioning assay, values from the following spindle dynamics metrics were used to compile the CDD: velocity, displacement per event, neck transit success rate, total spindle displacement per minute, and number of events per minute. The latter two were both included in the CDD tabulation since they revealed two different aspects of dynein activity: the extent to which the mutant dynein-dynactin complexes could move the spindle (total displacement per minute), and the ability to initiate a spindle movement event (number of events per minute). Since the spindle positioning assay reveals gross perturbations in dynein function, we increased the weight of this score with respect to each of the other metrics, such that the summed spindle dynamics metrics were weighed equally with that of the spindle positioning assay (see Figure 8, Supplement 1 for details on CDD tabulation). This analysis revealed a broad range of dynein dysfunction ranging from low (< 10) to high CDDs (> 30) for the mutant library. For comparison, the CDDs for wild-type and *dyn1*Δ cells were set to 0 and 100, respectively. Thus, considering the phenotypic severity of H3639P (CDD = 82) – which had a spindle positioning defect as severe as *dyn1*Δ, but exhibited some dynein activity in the spindle dynamics assays – we find that the CDD score accurately reflects the degree of dysfunction for each mutant.

Ranking the mutants according to their CDD scores (from low to high; Fig. 8B) revealed an apparent correlation between dynein dysfunction and disease type. Specifically, we found that above a CDD value of ~18, the likelihood of the mutation correlating with MCD increased, while its correlation with SMA-LED, CMT or CMD dropped. This suggests that the two general types of diseases (motor neuron disease, and defects in brain development) are each caused by different degrees of dynein dysfunction, and that there exists a lower threshold of dynein activity that is required for neurological development (see Discussion).

## DISCUSSION

Neurodegenerative or developmental diseases that arise as a consequence of mutations within the dynein gene – or dyneinopathies – are a broad range of devastating diseases that include muscular atrophy, muscular dystrophy, and malformations in cortical development (MCD). The ages of onset for these diseases range from birth to late adulthood, while the severity of the symptoms associated with them also cover a broad range^43–46^. Although the underlying reasons behind this symptomatic diversity are unclear, our findings suggest that the degree of dysfunction is at least a potential genetic determinant of the type of disease. Specifically, motor neuron diseases appear to be more susceptible to even small degrees of dysfunction (CDD from 5-18), while MCD tends to correlate with larger degrees of dynein dysfunction (CDD ≥ 19). The reasons for this are unclear, but we hypothesize that the differences are due to the somewhat distinct types of dynein-mediated transport required to maintain motor neuron health, versus that required to effect nuclear or neuronal migration in the developing neocortex. Motor neurons possess extremely long axons (≤ 1 m), and thus require a high degree of processive transport for the myriad vesicular cargoes that are moved from the soma to the axon terminal and back. This transport takes place not only during development and in growing children, but also in fully matured adults. Thus, it stands to reason that even subtle loss-of-functions in dynein-mediated transport can, over time, compromise motor neuron health. This is apparent from the wide range in ages-of-onset (from birth to adulthood), and in the range of severity for dynein-based motor neuron diseases (from weakness in the lower limbs to gross motor difficulties). On the other hand, during early brain development, dynein plays key roles in interkinetic nuclear migration (INM) and neuronal migration during development of the neocortex^30, 31, 89^. Nuclear envelope-anchored dyneins move the nucleus tens of microns from the basal to the apical surface of the neuroepithelium^31^. In addition to being a shorter distance traveled compared to axonal transport in a motor neuron, the number of motors engaged with microtubules during a nuclear migration event is likely far greater. Immunofluorescence reveals dynein is present along most of the nuclear envelope surface^31^. Moreover, live cell imaging revealed that the microtubule network is fairly extensive in the proximity of the nucleus^30, 89^. A similar process may be at play during post-INM neuronal migration, when dynein assists in centrosome advancement of the postmitotic neuron. Evidence suggests that, as in budding yeast, astral microtubules make contacts with teams of cortically anchored dynein motors to move the centrosome^89^. Thus, for both INM and neuronal migration, many dynein molecules are likely engaging with microtubules to effect neurogenesis. This is in contrast to a vesicle that is being transported along the axon of a motor neuron, which likely possesses far fewer motors (~3-7 per vesicle^11, 90^), of which potentially only a subset are engaged due to the geometric constraints associated with a three-dimensional vesicle engaging with a single filament. Thus, the minimal functional requirements for individual dynein motors during INM or neuronal migration are potentially lower due to the large number of motors engaged during a migration event. In such a scenario, teams of motors comprised of a mixture of wild-type and mutant variants (due to the heterozygous nature of these diseases) can work together to effect INM, but are less able to effectively transport single vesicular cargoes along the axon.

Our data indicate that K1475Q – a mutation in the linker domain – increases dynein run length *in vitro*, and alters its localization pattern *in vivo*. The position of this mutation (at an intermolecular interface that helps mediate formation of an auto-inhibited Phi particle conformation of human dynein^72^) suggests that the associated phenotypes may be a consequence of altered activity regulation. Given this mutant exhibited dynein-intrinsic enhancements in *in vitro* activity, the reduction in cellular dynein-dynactin activity is likely a consequence of the altered localization pattern. Specifically, the localization phenotype raises the possibility that a reduction in the number of dynein-dynactin complexes per cortical site (as apparent by the reduced fluorescence intensity of cortical foci; Fig. 5D) – which would result in fewer motor complexes being engaged for a spindle movement event – is the basis for cellular dysfunction. Given the auto-inhibited Phi particle exhibits reduced affinity for dynactin^72^, the altered localization pattern of the mutant may be due to disruption of a similar auto-inhibitory conformation in yeast, consequent increased dynactin binding, and thus an increase in the frequency of cortical off-loading events^63^. Previous studies have suggested that dynein’s interaction with dynactin is a limiting step in the delivery of dynein-dynactin complexes to cortical Num1 sites^77^. Future studies will be required to assess whether yeast dynein indeed adopts such an auto-inhibited state, and what role this conformation plays in the regulation of dynein activity.

In addition to revealing the potential molecular basis for disease onset or progression in affected patients, our findings also identified the potential structural basis for dynein dysfunction in two of the mutants. In the case of H3639P, our data support a model wherein the proline substitution compromises structural plasticity within an inter-helical loop in AAA5 that ultimately leads to destabilization and degradation of the protein. This conclusion is based on the reduced localization phenotype (Fig. 5B and C) that is rescued by flanking glycines (Fig. 6F) and proteasome inhibition (Fig. 6G), and the large proportion of non-motile microtubule-bound motors we observed in the single molecule assay (Fig. 4D). Since nearly all these motors exhibit persistent microtubule binding but no motility (96%; see kymograph in Figure 4, Supplement 1), we propose that a structural defect within the AAA ring is compromising the ability of nucleotide binding or hydrolysis to communicate with the microtubule-binding domain. It is interesting to note that the small fraction of motors that do exhibit processive motility *in vitro* (4.5%) move at velocities that are roughly similar to wild-type motors (80 versus 58 nm/sec for wild-type and H3639P, respectively). We observed a similar phenomenon during dynein-mediated spindle translocation *in vivo* (42 versus 22 nm/sec). These findings suggest that a small fraction of the motors are capable of overcoming folding defects to adopt a native, motility-competent conformation.

Subsequent to initiating this study, another group published findings describing the *in vitro* motility properties of a subset of the mutants analyzed here^71^. In this study, the authors utilized a recombinant human dynein in complex with dynactin and the adaptor BicD2 to assess single molecule motility parameters. With only two exceptions (E109I and N283R), the findings from this highly informative study largely corroborate our own data in that the dynein mutants assessed compromised the processivity of dynein-dynactin complexes (see Fig. 8A, left, “D”). For instance, the two mutants with the most severe phenotypic consequences were the same as those observed here (R1962C and H3822P). Although one of the exceptions – E109I – did not exhibit reduced processivity in our assays, the authors’ observation that this mutation leads to a reduction in the number of processive motors is similar to our observation of reduced cellular dynein-dynactin activity for this mutant (Fig. 8A, left, “A”).

In summary, we have established yeast as a medium-throughput model system that can be used to assess the molecular basis for dysfunction of disease-correlated dynein mutants. Our rapid and economical toolbox can be easily applied to understand the underlying basis for dysfunction in newly identified dynein mutants found in patients suffering from neurological diseases. We have also demonstrated the feasibility of rapidly testing hypotheses generated from our battery of assays using the wealth of available structural information that is available for dynein and its regulators.

## AUTHOR CONTRIBUTIONS

S.M.M. designed the study. M.G.M designed some of the experiments, performed most of them, and analyzed most of the data with support from J.M.G. Yeast strains were generated by M.G.M., J.M.G., K.M.R., and R.E.B. The Matlab code used to track the mitotic spindles was written by C.P.F. The manuscript was written by S.M.M. with assistance from M.G.M.

## ACKNOWLEDGEMENTS

We are grateful to Andrew Carter for sharing unpublished reagents (the human dynein complex expression plasmids) and valuable discussions. We also want to thank Richard Vallee, James Bamburg, Ashok Prasad, Olve Peersen, and members of the Markus and DeLuca laboratories for valuable discussions. This work was funded by the Muscular Dystrophy Association (376387 to S.M.M.) and the NIH/NIGMS (GM118492 to S.M.M.). We also thank Dr. Jeffrey Moore (and NIH R01-GM112893) for providing support for C.P.F. who developed Matlab code used for tracking spindles in live cells, and for sharing the *pdr1-DBD-CYC8* yeast strain.

## METHODS

### Media and strain construction

Strains are derived from either W303 or YEF473A^92^ and are listed in Table S1. We transformed yeast strains using the lithium acetate method^93^. Strains carrying mutations were constructed by PCR product-mediated transformation^94^ or by mating followed by tetrad dissection. Proper tagging and mutagenesis was confirmed by PCR, and in most cases sequencing (all point mutations were confirmed via sequencing). Fluorescent tubulin-expressing yeast strains were generated using plasmids and strategies described previously^95, 96^. Yeast synthetic defined (SD) media was obtained from Sunrise Science Products (San Diego, CA).

### Plasmid generation

For expression and purification of human dynein complex mutants (or wild-type), mutations were engineered into the human dynein heavy chain (DHC)-containing plasmid, pbiG1a:6His-ZZ-SNAPf-DHC1. We used Gibson assembly to engineer point mutations – C1932S, R1962C, or both – into this plasmid. The PmeI-digested gene expression cassette from this plasmid was co-assembled with the PmeI-digested poly-gene cassette from pbiG1b:IC2/LIC2/Tctex1/Robl1/LC8 (encoding all dynein accessory chains) into PmeI-digested pbiG2ab using biGBac cloning strategies as previously described^97^ (all wild-type plasmids were kind gifts from Andrew Carter). The final, sequence-verified plasmids (wild-type and mutant variants of pbiG2ab:6His-ZZ-SNAPf-DHC1/IC2/LIC2/Tctex1/Robl1/LC8) were used to generate recombinant baculoviral genomes by Tn7 transposition into DH10Bac cells (Life Technologies). White, PCR-confirmed colonies were inoculated into LB media supplemented with 7 µg/ml gentamycin, 10 µg/ml tetracycline and 50 µg/ml kanamycin and grown overnight at 37°C. Bacmid preparation was performed as described previously^72^, stored at 4°C, and used within 2 weeks for subsequent virus production (see below).

### Protein purification

Purification of yeast dynein (ZZ-TEV-Dyn1-HALO, under the native *DYN1* promoter; or, ZZ-TEV-6His-GFP-3HA-GST-dynein_331_-HALO, under the control of the galactose-inducible promoter, *GAL1p*) was performed as previously described ^70, 98^. Briefly, yeast cultures were grown in YPA supplemented with either 2% glucose (for full-length dynein) or 2% galactose (for GST-dynein_331_), harvested, washed with cold water, and then resuspended in a small volume of water. The resuspended cell pellet was drop frozen into liquid nitrogen and then lysed in a coffee grinder (Hamilton Beach). After lysis, 0.25 volume of 4X dynein lysis buffer (1X buffer: 30 mM HEPES, pH 7.2, 50 mM potassium acetate, 2 mM magnesium acetate, 0.2 mM EGTA) supplemented with 1 mM DTT, 0.1 mM Mg-ATP, 0.5 mM Pefabloc SC (concentrations for 1X buffer) was added, and the lysate was clarified at 22,000 × g for 20 min. The supernatant was then bound to IgG sepharose 6 fast flow resin (GE) for 1-1.5 hours at 4°C, which was subsequently washed three times in 5 ml lysis buffer, and twice in TEV buffer (50 mM Tris, pH 8.0, 150 mM potassium acetate, 2 mM magnesium acetate, 1 mM EGTA, 0.005% Triton X-100, 10% glycerol, 1 mM DTT, 0.1 mM Mg-ATP, 0.5 mM Pefabloc SC). To fluorescently label the motors for single molecule analyses, the bead-bound protein was incubated with either 6.7 µM HaloTag-AlexaFluor660 (Promega) for 10 minutes at room temperature. The resin was then washed four more times in TEV digest buffer, then incubated in TEV buffer supplemented with TEV protease for 1-1.5 hours at 16°C. Following TEV digest, the beads were pelleted, and the resulting supernatant was aliquoted, flash frozen in liquid nitrogen, and stored at −80°C.

The human dynein complex was expressed and purified from insect cells (ExpiSf9 cells; Life Technologies) as previously described with minor modifications^72, 83^. Briefly, 4 ml of ExpiSf9 cells at 2.5 × 10^6^ cells/ml, which were maintained in ExpiSf CD Medium (Life Technologies), were transfected with 1 µg of bacmid DNA (see above) using ExpiFectamine (Life Technologies) according to the manufacturer’s instructions. 5 days following transfection, the cells were pelleted, and 1 ml of the resulting supernatant (P1) was used to infect 300 ml of ExpiSf9 cells (5 × 10^6^ cells/ml). 72 hours later, the cells were harvested (2000 × g, 20 min), washed with phosphate buffered saline (pH 7.2), pelleted again (1810 × g, 20 min), and resuspended in an equal volume of human dynein lysis buffer (50 mM HEPES, pH 7.4, 100 mM NaCl, 10% glycerol, 1 mM DTT, 0.1 mM Mg-ATP, 1 mM PMSF). The resulting cell suspension was drop frozen in liquid nitrogen and stored at −80°C. For protein purification, 30 ml of additional human dynein lysis buffer supplemented with cOmplete protease inhibitor cocktail (Roche) was added to the frozen cell pellet, which was then rapidly thawed in a 37°C water bath prior to incubation on ice. Cells were lysed in a dounce-type tissue grinder (Wheaton) using ≥ 150 strokes (lysis was monitored by microscopy). Subsequent to clarification at 40,000 × g, 45 min, the supernatant was applied to 2 ml of IgG sepharose fast flow resin (GE) pre-equilibrated in human dynein lysis buffer, and incubated at 4°C for 2-4 hours. Beads were then washed with 50 ml of human dynein lysis buffer, and 50 ml of human dynein TEV buffer (50 mM Tris pH 7.4, 150 mM potassium acetate, 2 mM magnesium acetate, 1 mM EGTA, 10% glycerol, 1 mM DTT, 0.1 mM Mg-ATP). The bead-bound protein was incubated with 3 µM SNAP-Surface Alexa Fluor 647 (NEB) for 40-60 min at 4°C (to fluorescently label the protein), washed 5 times in human dynein TEV buffer, then incubated with TEV protease overnight at 4°C. The next morning, the recovered supernatant was applied to a Superose 6 gel filtration column (GE) equilibrated in GF150 buffer (25 mM HEPES pH 7.4, 150 mM KCl, 1 mM MgCl_2_, 5 mM DTT, 0.1 mM Mg-ATP) using an AKTA Pure. Peak fractions (determined by UV 260 nm absorbance and SDS-PAGE) were pooled, concentrated, aliquoted, flash frozen, then stored at −80°C.

### Single and ensemble molecule motility assays

The yeast dynein single-molecule motility assay was performed as previously described with minor modifications^70^. Briefly, flow chambers constructed using slides and plasma cleaned and silanized coverslips attached with double-sided adhesive tape were coated with anti-tubulin antibody (8 μg/ml, YL1/2; Accurate Chemical & Scientific Corporation) then blocked with 1% Pluronic F-127 (Fisher Scientific). Taxol-stabilized microtubules assembled from unlabeled and fluorescently-labeled porcine tubulin (10:1 ratio; Cytoskeleton) were introduced into the chamber. Following a 5-10 minute incubation, the chamber was washed with dynein lysis buffer (see above) supplemented with 20 μM taxol, and then purified dynein motors were introduced in the chamber. After a 1 minute incubation, motility buffer (30 mM HEPES pH 7.2, 50 mM potassium acetate, 2 mM magnesium acetate, 1 mM EGTA, 1 mM DTT, 1 mM Mg-ATP) supplemented with 0.05% Pluronic F-127, 20 µM taxol, and an oxygen-scavenging system (1.5% glucose, 1 U/ml glucose oxidase, 125 U/ml catalase) was added. TIRFM images were collected using a 1.49 NA 100X TIRF objective on a Nikon Ti-E inverted microscope equipped with a Ti-S-E motorized stage, piezo Z-control (Physik Instrumente), and an iXon X3 DU897 cooled EM-CCD camera (Andor). 488 nm, 561 nm, and 640 nm lasers (Coherent) were used along with a multi-pass quad filter cube set (C-TIRF for 405/488/561/638 nm; Chroma) and emission filters mounted in a filter wheel (525/50 nm, 600/50 nm and 700/75 nm; Chroma). We acquired images at 2 second intervals for 8 min. Velocity and run length values were determined from kymographs generated using the MultipleKymograph plugin for ImageJ (http://www.embl.de/eamnet/html/body_kymograph.html).

Human dynein-mediated microtubule gliding assays were performed as previously described^72^ with minor modifications. Briefly, flow chambers were prepared by affixing an ethanol-flamed coverslip to a glass slide using double-stick tape. The chamber was then incubated on an ice block, washed with 1% Pluronic F-127, following by addition of purified dynein (5 chamber volumes of 60 nM dynein complex). Unbound motors were washed out with GF150 buffer. Subsequently, motility buffer (30 mM HEPES pH 7.0, 50 mM KCl, 5 mM MgSO4, 1 mM EGTA, 1 mM DTT, 2.5 mM Mg-ATP, 40 µM taxol) supplemented with 1.5% glucose, the oxygen scavenging system (see above), and 150 nM fluorescent microtubules was added to the chamber. Images were acquired every 1 sec (for wild-type) or 5 sec (for mutants), and velocity values were determined from kymographs generated as above.

### Live cell imaging experiments

For the single time point spindle position assay, the percentage of cells with a misoriented anaphase spindle was determined after growth overnight (12-16 hours) at a low temperature (16°C), as previously described^78, 88, 99^. A single z-stack of wide-field fluorescence images was acquired for mRuby2-Tub1. For the spindle dynamics assay, cells were arrested with hydroxyurea (HU) for 2.5 hours, and then mounted on agarose pads containing HU for fluorescence microscopy. Full Z-stacks (23 planes at 0.2 µm spacing) of GFP-labeled microtubules (GFP-Tub1) were acquired every 10 seconds for 10 minutes on a stage pre-warmed to 30°C. To image dynein localization in live cells, cells were grown to mid-log phase in SD media supplemented with 2% glucose, and mounted on agarose pads. Images were collected on a Nikon Ti-E microscope equipped with a 1.49 NA 100X TIRF objective, a Ti-S-E motorized stage, piezo Z-control (Physik Instrumente), an iXon DU888 cooled EM-CCD camera (Andor), a stage-top incubation system (Okolab), and a spinning disc confocal scanner unit (CSUX1; Yokogawa) with an emission filter wheel (ET525/50M for GFP, and ET632/60M for mRuby2; Chroma). Lasers (488 nm and 561 nm) housed in a LU-NV laser unit equipped with AOTF control (Nikon) were used to excite GFP and mRuby2, respectively. The microscope was controlled with NIS Elements software (Nikon).

### Statistical analyses

Statistical tests were performed as described in the figure legends. Unpaired Welch’s t tests (for gaussian distributed velocity data) and Mann-Whitney test (for exponentially distributed dispalcement data) were performed using Graphpad Prism. Z scores, which are a quantitative measure of difference between two proportions, were calculated using the following formula:

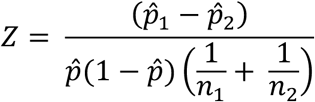

where:

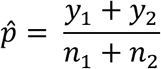

Z scores were converted to two-tailed P values using an online calculator.

### Coefficient of dynein dysfunction (CDD) score calculation

To calculate the CDD scores, we used the following approach which permitted a quantitative measure of difference between mean values obtained for wild-type versus those obtained for each mutant. Graphpad Prism was used to calculate q values (*i.e.*, the difference between the two means divided by the standard error of that difference), whereas Z scores were calculated as described above (all values are shown in Figure 8, Supplement 1). We then converted the q values and Z scores for each mutant (for each assay) into a “normalized relative variance” score (nrv), which reflects the relative difference between two mean values (*e.g.*, between wild-type and mutant 1; as reflected in the Z scores and q values, or “v”), where nrv = |v|/v_max_ for each range of scores (for each column shown in Figure 8, Supplement 1A). To convert the nrv values into a final CDD score for each mutant, we used the formula shown in Figure 8, Supplement 1C. Briefly, the nrv values for each assay for a given mutant was added, with the spindle positioning nrv (nrv_SP_) weighed five times that of the others, as described within the Results. In the two cases where a value wasn’t determined (due to insufficient observations, such as in the case for neck transit success for the H3639P mutant), the denominator was reduced from 6 to 5. In the two instances where the Z score for spindle positioning was negative (due to a lower number of mispositioned spindles being observed in K540C and D2439K cells than in wild-type cells; see Fig. 1B), we adjusted the values to 0 so as to avoid them skewing the nrv_SP_ values.

## REFERENCES

1. Roberts, A. J., Kon, T., Knight, P. J., Sutoh, K. & Burgess, S. A. in Nat Rev Mol Cell Biol 1–14 (Nature Publishing Group, 2013).

2. Vallee, R. B., Mckenney, R. J. & Ori-Mckenney, K. M. in Nature cell Biology Vol. 14 224–230 (Nature Publishing Group, 2012).

3. Kardon, J. R. & Vale, R. D. Regulators of the cytoplasmic dynein motor. Nat Rev Mol Cell Biol 10, 854–865, (2009).

4. Xia, C. H. et al. Abnormal neurofilament transport caused by targeted disruption of neuronal kinesin heavy chain KIF5A. J Cell Biol 161, 55–66, (2003).

5. He, Y. et al. Role of cytoplasmic dynein in the axonal transport of microtubules and neurofilaments. J Cell Biol 168, 697–703, (2005).

6. Wagner, O. I. et al. The interaction of neurofilaments with the microtubule motor cytoplasmic dynein. Mol Biol Cell 15, 5092–5100, (2004).

7. Shah, J. V., Flanagan, L. A., Janmey, P. A. & Leterrier, J. F. Bidirectional translocation of neurofilaments along microtubules mediated in part by dynein/dynactin. Mol Biol Cell 11, 3495–3508, (2000).

8. Bowman, A. B. et al. Kinesin-dependent axonal transport is mediated by the sunday driver (SYD) protein. Cell 103, 583–594, (2000).

9. Berg, E. A. et al. Isolation and characterization of substance P-containing dense core vesicles from rabbit optic nerve and termini. J Neurosci Res 62, 830–839, (2000).

10. Barkus, R. V., Klyachko, O., Horiuchi, D., Dickson, B. J. & Saxton, W. M. Identification of an axonal kinesin-3 motor for fast anterograde vesicle transport that facilitates retrograde transport of neuropeptides. Mol Biol Cell 19, 274–283, (2008).

11. Hendricks, A. G. et al. Motor coordination via a tug-of-war mechanism drives bidirectional vesicle transport. Curr Biol 20, 697–702, (2010).

12. Rosa-Ferreira, C. & Munro, S. Arl8 and SKIP act together to link lysosomes to kinesin-1. Dev Cell 21, 1171–1178, (2011).

13. Maday, S., Wallace, K. E. & Holzbaur, E. L. Autophagosomes initiate distally and mature during transport toward the cell soma in primary neurons. J Cell Biol 196, 407–417, (2012).

14. Kamal, A., Stokin, G. B., Yang, Z., Xia, C. H. & Goldstein, L. S. Axonal transport of amyloid precursor protein is mediated by direct binding to the kinesin light chain subunit of kinesin-I. Neuron 28, 449–459, (2000).

15. Lazarov, O. et al. Axonal transport, amyloid precursor protein, kinesin-1, and the processing apparatus: revisited. J Neurosci 25, 2386–2395, (2005).

16. Fu, M. M. & Holzbaur, E. L. JIP1 regulates the directionality of APP axonal transport by coordinating kinesin and dynein motors. J Cell Biol 202, 495–508, (2013).

17. Almenar-Queralt, A. et al. UV irradiation accelerates amyloid precursor protein (APP) processing and disrupts APP axonal transport. J Neurosci 34, 3320–3339, (2014).

18. Rao, A. N. et al. Cytoplasmic Dynein Transports Axonal Microtubules in a Polarity-Sorting Manner. Cell Rep 19, 2210–2219, (2017).

19. Ravikumar, B. et al. Dynein mutations impair autophagic clearance of aggregate-prone proteins. Nature genetics 37, 771–776, (2005).

20. Katsumata, K. et al. Dynein- and activity-dependent retrograde transport of autophagosomes in neuronal axons. Autophagy 6, 378–385, (2010).

21. Cheng, X. T., Zhou, B., Lin, M. Y., Cai, Q. & Sheng, Z. H. Axonal autophagosomes recruit dynein for retrograde transport through fusion with late endosomes. J Cell Biol 209, 377–386, (2015).

22. Horak, M., Petralia, R. S., Kaniakova, M. & Sans, N. ER to synapse trafficking of NMDA receptors. Front Cell Neurosci 8, 394, (2014).

23. Levy, J. R. et al. A motor neuron disease-associated mutation in p150Glued perturbs dynactin function and induces protein aggregation. J Cell Biol 172, 733–745, (2006).

24. LaMonte, B. H. et al. Disruption of dynein/dynactin inhibits axonal transport in motor neurons causing late-onset progressive degeneration. Neuron 34, 715–727, (2002).

25. Lin, H. & Schlaepfer, W. W. Role of neurofilament aggregation in motor neuron disease. Ann Neurol 60, 399–406, (2006).

26. Schiavo, G., Greensmith, L., Hafezparast, M. & Fisher, E. M. Cytoplasmic dynein heavy chain: the servant of many masters. Trends Neurosci 36, 641–651, (2013).

27. Hafezparast, M. et al. Mutations in dynein link motor neuron degeneration to defects in retrograde transport. Science 300, 808–812, (2003).

28. Chen, X. J. et al. Proprioceptive sensory neuropathy in mice with a mutation in the cytoplasmic Dynein heavy chain 1 gene. J Neurosci 27, 14515–14524, (2007).

29. Ori-McKenney, K. M., Xu, J., Gross, S. P. & Vallee, R. B. A cytoplasmic dynein tail mutation impairs motor processivity. Nat Cell Biol 12, 1228–1234, (2010).

30. Tsai, J. W., Lian, W. N., Kemal, S., Kriegstein, A. R. & Vallee, R. B. Kinesin 3 and cytoplasmic dynein mediate interkinetic nuclear migration in neural stem cells. Nat Neurosci 13, 1463–1471, (2010).

31. Hu, D. J. et al. Dynein recruitment to nuclear pores activates apical nuclear migration and mitotic entry in brain progenitor cells. Cell 154, 1300–1313, (2013).

32. Del Bene, F., Wehman, A. M., Link, B. A. & Baier, H. Regulation of neurogenesis by interkinetic nuclear migration through an apical-basal notch gradient. Cell 134, 1055–1065, (2008).

33. Harms, M. B. et al. Mutations in the tail domain of DYNC1H1 cause dominant spinal muscular atrophy. Neurology 78, 1714–1720, (2012).

34. Tsurusaki, Y. et al. A DYNC1H1 mutation causes a dominant spinal muscular atrophy with lower extremity predominance. Neurogenetics 13, 327–332, (2012).

35. Scoto, M. et al. Novel mutations expand the clinical spectrum of DYNC1H1-associated spinal muscular atrophy. Neurology, (2015).

36. Peeters, K. et al. Novel mutations in the DYNC1H1 tail domain refine the genetic and clinical spectrum of dyneinopathies. Human mutation, (2014).

37. Strickland, A. V. et al. Mutation screen reveals novel variants and expands the phenotypes associated with DYNC1H1. J Neurol 262, 2124–2134, (2015).

38. Weedon, M. N. et al. Exome sequencing identifies a DYNC1H1 mutation in a large pedigree with dominant axonal Charcot-Marie-Tooth disease. American journal of human genetics 89, 308–312, (2011).

39. Fiorillo, C. et al. Novel dynein DYNC1H1 neck and motor domain mutations link distal spinal muscular atrophy and abnormal cortical development. Human mutation 35, 298–302, (2014).

40. Wynshaw-Boris, A. Lissencephaly and LIS1: insights into the molecular mechanisms of neuronal migration and development. Clin Genet 72, 296–304, (2007).

41. Wynshaw-Boris, A. & Gambello, M. J. LIS1 and dynein motor function in neuronal migration and development. Genes & development 15, 639–651, (2001).

42. Reiner, O. et al. Isolation of a Miller-Dieker lissencephaly gene containing G protein beta-subunit-like repeats. Nature 364, 717–721, (1993).

43. Poirier, K. et al. Mutations in TUBG1, DYNC1H1, KIF5C and KIF2A cause malformations of cortical development and microcephaly. Nature genetics 45, 639–647, (2013).

44. Laquerriere, A. et al. Neuropathological Hallmarks of Brain Malformations in Extreme Phenotypes Related to DYNC1H1 Mutations. J Neuropathol Exp Neurol 76, 195–205, (2017).

45. Willemsen, M. H. et al. Mutations in DYNC1H1 cause severe intellectual disability with neuronal migration defects. Journal of medical genetics 49, 179–183, (2012).

46. Vissers, L. E. et al. A de novo paradigm for mental retardation. Nature genetics 42, 1109–1112, (2010).

47. Rusan, N. M., Tulu, U. S., Fagerstrom, C. & Wadsworth, P. Reorganization of the microtubule array in prophase/prometaphase requires cytoplasmic dynein-dependent microtubule transport. J Cell Biol 158, 997–1003, (2002).

48. Gonczy, P., Pichler, S., Kirkham, M. & Hyman, A. A. Cytoplasmic dynein is required for distinct aspects of MTOC positioning, including centrosome separation, in the one cell stage Caenorhabditis elegans embryo. J Cell Biol 147, 135–150, (1999).

49. Salina, D. et al. Cytoplasmic dynein as a facilitator of nuclear envelope breakdown. Cell 108, 97–107, (2002).

50. Howell, B. J. et al. Cytoplasmic dynein/dynactin drives kinetochore protein transport to the spindle poles and has a role in mitotic spindle checkpoint inactivation. J Cell Biol 155, 1159–1172, (2001).

51. Wojcik, E. et al. Kinetochore dynein: its dynamics and role in the transport of the Rough deal checkpoint protein. Nat Cell Biol 3, 1001–1007, (2001).

52. Chevalier-Larsen, E. & Holzbaur, E. L. Axonal transport and neurodegenerative disease. Biochim Biophys Acta 1762, 1094–1108, (2006).

53. Li, Y. Y., Yeh, E., Hays, T. & Bloom, K. Disruption of mitotic spindle orientation in a yeast dynein mutant. Proc Natl Acad Sci U S A 90, 10096–10100, (1993).

54. Eshel, D. et al. Cytoplasmic dynein is required for normal nuclear segregation in yeast. Proc Natl Acad Sci U S A 90, 11172–11176, (1993).

55. Carminati, J. L. & Stearns, T. Microtubules orient the mitotic spindle in yeast through dynein-dependent interactions with the cell cortex. J Cell Biol 138, 629–641, (1997).

56. Markus, S. M. & Lee, W. L. Microtubule-dependent path to the cell cortex for cytoplasmic dynein in mitotic spindle orientation. Bioarchitecture 1, 209–215, (2011).

57. Pfister, K. K. et al. Genetic analysis of the cytoplasmic dynein subunit families. PLoS Genet 2, e1, (2006).

58. Raaijmakers, J. A., Tanenbaum, M. E. & Medema, R. H. Systematic dissection of dynein regulators in mitosis. J Cell Biol 201, 201–215, (2013).

59. Carter, A. P. Crystal clear insights into how the dynein motor moves. J Cell Sci 126, 705–713, (2013).

60. Schmidt, H. & Carter, A. P. Review: Structure and mechanism of the dynein motor ATPase. Biopolymers 105, 557–567, (2016).

61. Reck-Peterson, S. L. et al. Single-molecule analysis of dynein processivity and stepping behavior. Cell 126, 335–348, (2006).

62. Markus, S. M., Kalutkiewicz, K. A. & Lee, W. L. She1-mediated inhibition of dynein motility along astral microtubules promotes polarized spindle movements. Curr Biol 22, 2221–2230, (2012).

63. Markus, S. M. & Lee, W. L. Regulated offloading of cytoplasmic dynein from microtubule plus ends to the cortex. Dev Cell 20, 639–651, (2011).

64. Urnavicius, L. et al. The structure of the dynactin complex and its interaction with dynein. Science 347, 1441–1446, (2015).

65. Schmidt, H., Zalyte, R., Urnavicius, L. & Carter, A. P. Structure of human cytoplasmic dynein-2 primed for its power stroke. Nature 518, 435–438, (2015).

66. Yin, H., Pruyne, D., Huffaker, T. C. & Bretscher, A. Myosin V orientates the mitotic spindle in yeast. Nature 406, 1013–1015, (2000).

67. Hwang, E., Kusch, J., Barral, Y. & Huffaker, T. C. Spindle orientation in Saccharomyces cerevisiae depends on the transport of microtubule ends along polarized actin cables. J Cell Biol 161, 483–488, (2003).

68. Liakopoulos, D., Kusch, J., Grava, S., Vogel, J. & Barral, Y. Asymmetric loading of Kar9 onto spindle poles and microtubules ensures proper spindle alignment. Cell 112, 561–574, (2003).

69. Moore, J. K., Sept, D. & Cooper, J. A. Neurodegeneration mutations in dynactin impair dynein-dependent nuclear migration. Proc Natl Acad Sci U S A 106, 5147–5152, (2009).

70. Ecklund, K. H. et al. She1 affects dynein through direct interactions with the microtubule and the dynein microtubule-binding domain. Nature communications 8, 2151, (2017).

71. Hoang, H. T., Schlager, M. A., Carter, A. P. & Bullock, S. L. DYNC1H1 mutations associated with neurological diseases compromise processivity of dynein-dynactin-cargo adaptor complexes. Proc Natl Acad Sci U S A 114, E1597–E1606, (2017).

72. Zhang, K. et al. Cryo-EM Reveals How Human Cytoplasmic Dynein Is Auto-inhibited and Activated. Cell 169, 1303–1314 e1318, (2017).

73. Amos, L. A. Brain dynein crossbridges microtubules into bundles. J Cell Sci 93 (Pt 1), 19–28, (1989).

74. Urnavicius, L. et al. Cryo-EM shows how dynactin recruits two dyneins for faster movement. Nature 554, 202–206, (2018).

75. Natan, E., Wells, J. N., Teichmann, S. A. & Marsh, J. A. Regulation, evolution and consequences of cotranslational protein complex assembly. Curr Opin Struct Biol 42, 90–97, (2017).

76. Nicholls, C. D., McLure, K. G., Shields, M. A. & Lee, P. W. Biogenesis of p53 involves cotranslational dimerization of monomers and posttranslational dimerization of dimers. Implications on the dominant negative effect. J Biol Chem 277, 12937–12945, (2002).

77. Markus, S. M. et al. Quantitative analysis of Pac1/LIS1-mediated dynein targeting: Implications for regulation of dynein activity in budding yeast. Cytoskeleton (Hoboken) 68, 157–174, (2011).

78. Li, J., Lee, W. L. & Cooper, J. A. NudEL targets dynein to microtubule ends through LIS1. Nat Cell Biol 7, 686–690, (2005).

79. Lammers, L. G. & Markus, S. M. The dynein cortical anchor Num1 activates dynein motility by relieving Pac1/LIS1-mediated inhibition. J Cell Biol 211, 309–322, (2015).

80. Lee, W. L., Oberle, J. R. & Cooper, J. A. The role of the lissencephaly protein Pac1 during nuclear migration in budding yeast. J Cell Biol 160, 355–364, (2003).

81. Heil-Chapdelaine, R. A., Oberle, J. R. & Cooper, J. A. The cortical protein Num1p is essential for dynein-dependent interactions of microtubules with the cortex. J Cell Biol 151, 1337–1344, (2000).

82. McKenney, R. J., Huynh, W., Tanenbaum, M. E., Bhabha, G. & Vale, R. D. Activation of cytoplasmic dynein motility by dynactin-cargo adapter complexes. Science 345, 337–341, (2014).

83. Schlager, M. A., Hoang, H. T., Urnavicius, L., Bullock, S. L. & Carter, A. P. In vitro reconstitution of a highly processive recombinant human dynein complex. EMBO J 33, 1855–1868, (2014).

84. Derr, N. D. et al. Tug-of-war in motor protein ensembles revealed with a programmable DNA origami scaffold. Science 338, 662–665, (2012).

85. Huynh, W. & Vale, R. D. Disease-associated mutations in human BICD2 hyperactivate motility of dynein-dynactin. J Cell Biol 216, 3051–3060, (2017).

86. Moore, J. K., Li, J. & Cooper, J. A. Dynactin function in mitotic spindle positioning. Traffic 9, 510–527, (2008).

87. Lee, W. L., Kaiser, M. A. & Cooper, J. A. The offloading model for dynein function: differential function of motor subunits. J Cell Biol 168, 201–207, (2005).

88. Markus, S. M., Punch, J. J. & Lee, W. L. Motor- and tail-dependent targeting of dynein to microtubule plus ends and the cell cortex. Curr Biol 19, 196–205, (2009).

89. Tsai, J. W., Bremner, K. H. & Vallee, R. B. Dual subcellular roles for LIS1 and dynein in radial neuronal migration in live brain tissue. Nat Neurosci 10, 970–979, (2007).

90. Rai, A. K., Rai, A., Ramaiya, A. J., Jha, R. & Mallik, R. Molecular adaptations allow dynein to generate large collective forces inside cells. Cell 152, 172–182, (2013).

91. Stepanov, A., Nitiss, K. C., Neale, G. & Nitiss, J. L. Enhancing drug accumulation in Saccharomyces cerevisiae by repression of pleiotropic drug resistance genes with chimeric transcription repressors. Mol Pharmacol 74, 423–431, (2008).

92. Bi, E. & Pringle, J. R. ZDS1 and ZDS2, genes whose products may regulate Cdc42p in Saccharomyces cerevisiae. Mol Cell Biol 16, 5264–5275, (1996).

93. Knop, M. et al. Epitope tagging of yeast genes using a PCR-based strategy: more tags and improved practical routines. Yeast 15, 963–972, (1999).

94. Longtine, M. S. et al. Additional modules for versatile and economical PCR-based gene deletion and modification in Saccharomyces cerevisiae. Yeast 14, 953–961, (1998).

95. Markus, S. M., Omer, S., Baranowski, K. & Lee, W. L. Improved Plasmids for Fluorescent Protein Tagging of Microtubules in Saccharomyces cerevisiae. Traffic 16, 773–786, (2015).

96. Song, S. & Lee, K. S. A novel function of Saccharomyces cerevisiae CDC5 in cytokinesis. J Cell Biol 152, 451–469, (2001).

97. Weissmann, F. et al. biGBac enables rapid gene assembly for the expression of large multisubunit protein complexes. Proc Natl Acad Sci U S A 113, E2564–2569, (2016).

98. Huang, J., Roberts, A. J., Leschziner, A. E. & Reck-Peterson, S. L. Lis1 acts as a “clutch” between the ATPase and microtubule-binding domains of the dynein motor. Cell 150, 975–986, (2012).

99. Sheeman, B. et al. Determinants of S. cerevisiae dynein localization and activation: implications for the mechanism of spindle positioning. Curr Biol 13, 364–372, (2003).

