## Supplemental Information for "Molecular basis for dyneinopathies reveals insight into dynein regulation and dysfunction"

**SUPPLEMENTARY INFORMATION**
**SUPPLEMENTARY FIGURES AND TABLE**

#### Figure 2, Supplement 1

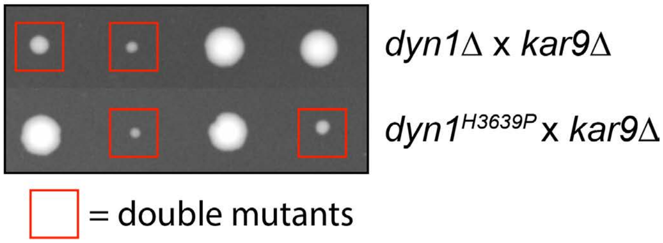

Figure 2, Supplement 1. **H3639P mutant exhibits synthetic genetic interactions with *KAR9*.** Cells expressing dyn1$^{H3639P}$ exhibit synthetic growth defects with *kar9*Δ that are as severe as *dyn1*Δ *kar9*Δ. Tetrads were dissected on YPAD media, and subsequently genotyped by growth on selective media. One representative tetrad each from a mating of a *kar9*Δ strain with either *dyn1*$^{H3639P}$ or *dyn1*Δ is shown. Double mutants are indicated with red boxes. Note that none of the other dynein mutants exhibited apparent synthetic growth defects with *kar9*Δ (not shown).

### Figure 2, Supplement 2

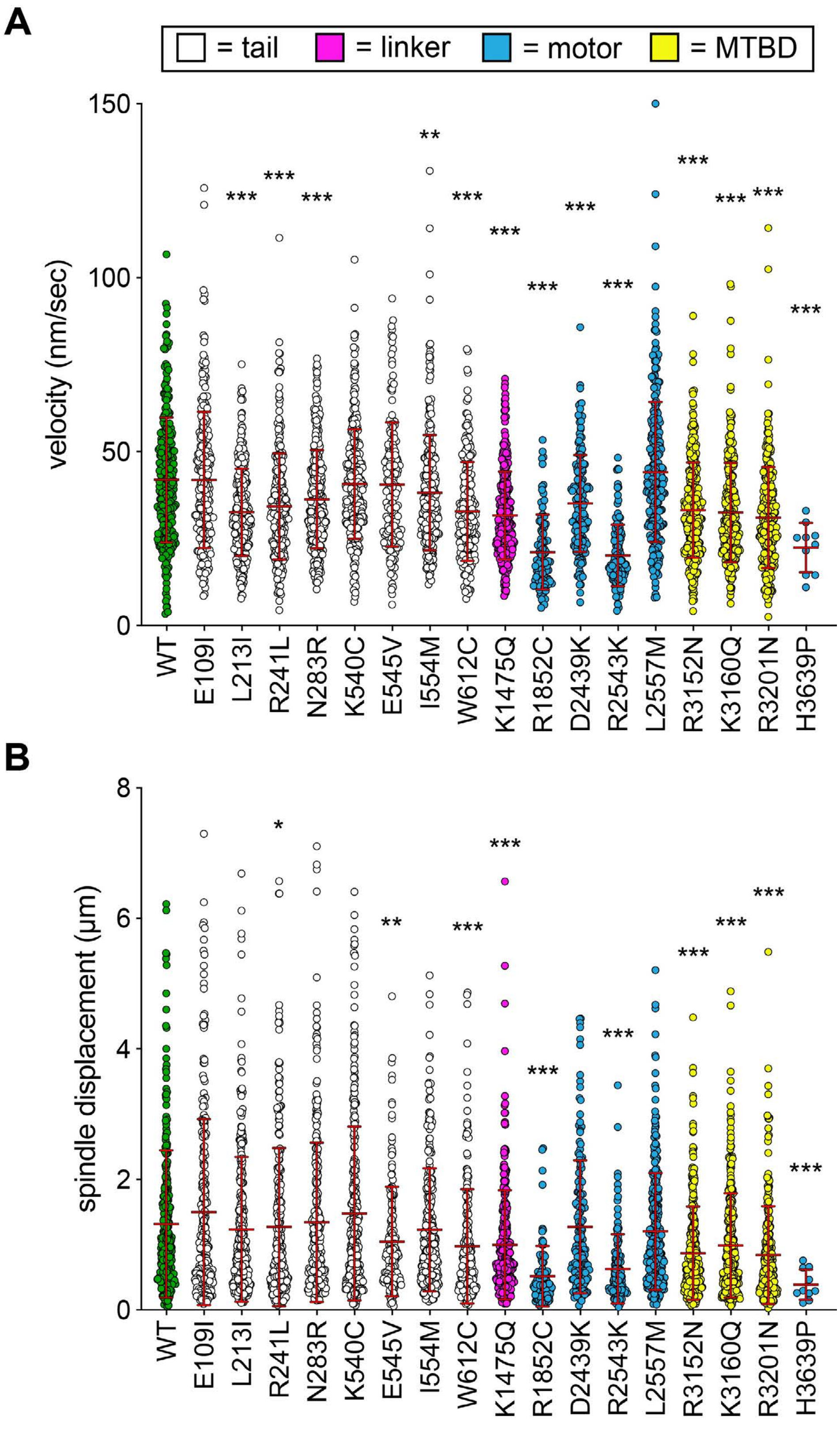

Figure 2, Supplement 2. **Additional plots of spindle dynamics data from haploid cells.** (A and B) Scatter plots depicting (A) velocity per spindle displacement event, and (B) displacement per spindle displacement event (from Figure 2). Mean values and standard deviations are also depicted with red lines.

**Figure 2, Supplement 3**

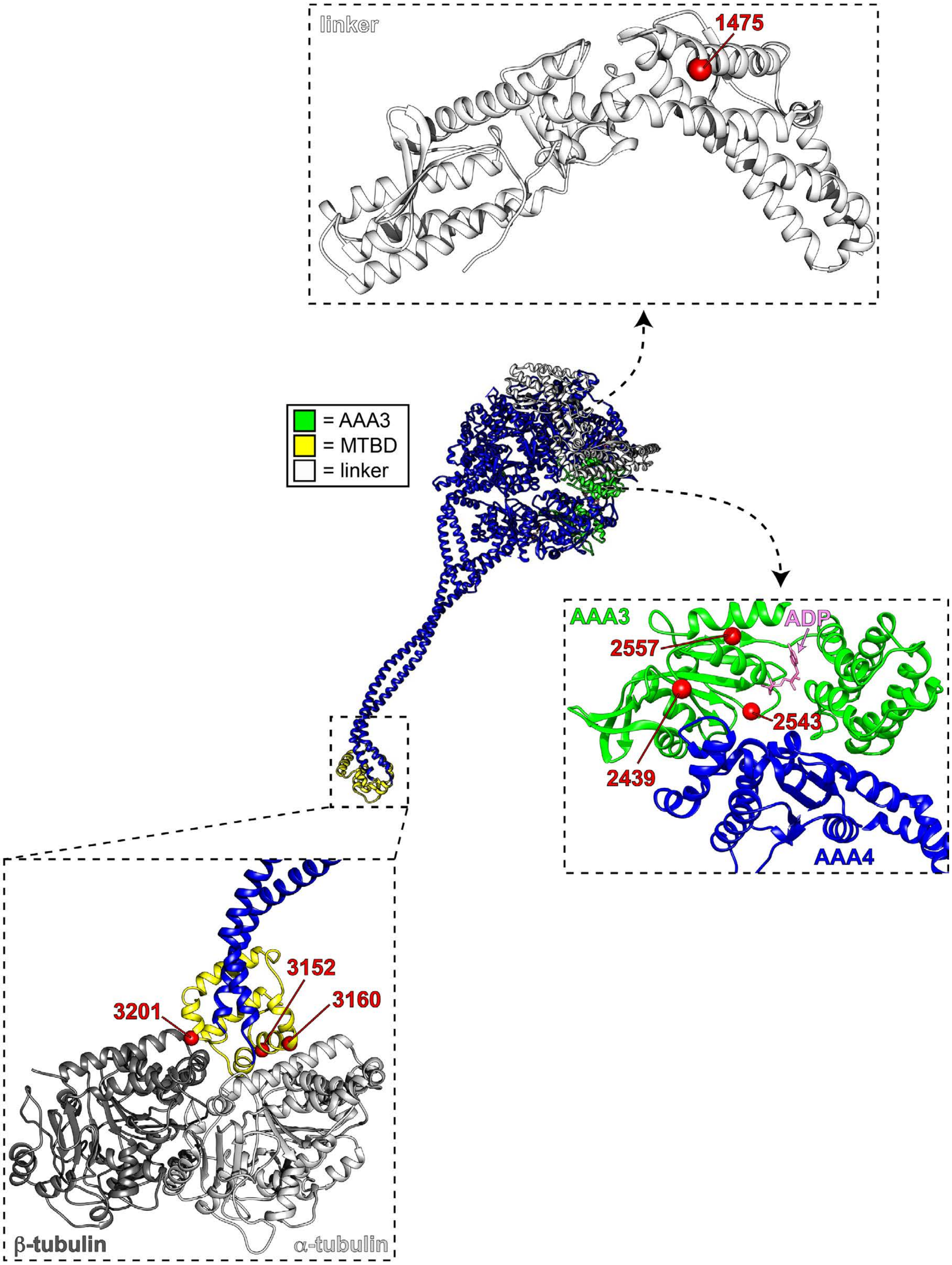

Figure 2, Supplement 3. **Mutations map to various structural elements within the motor domain.** Structural model of the dynein motor domain (from PDB 4RH7[1]) with zoomed in regions depicting the linker domain, the AAA3/AAA4 interface, and the MTBD bound to microtubules (R4H7 docked into 3J1T[2]). Red spheres denote mutations.

### Figure 2, Supplement 4

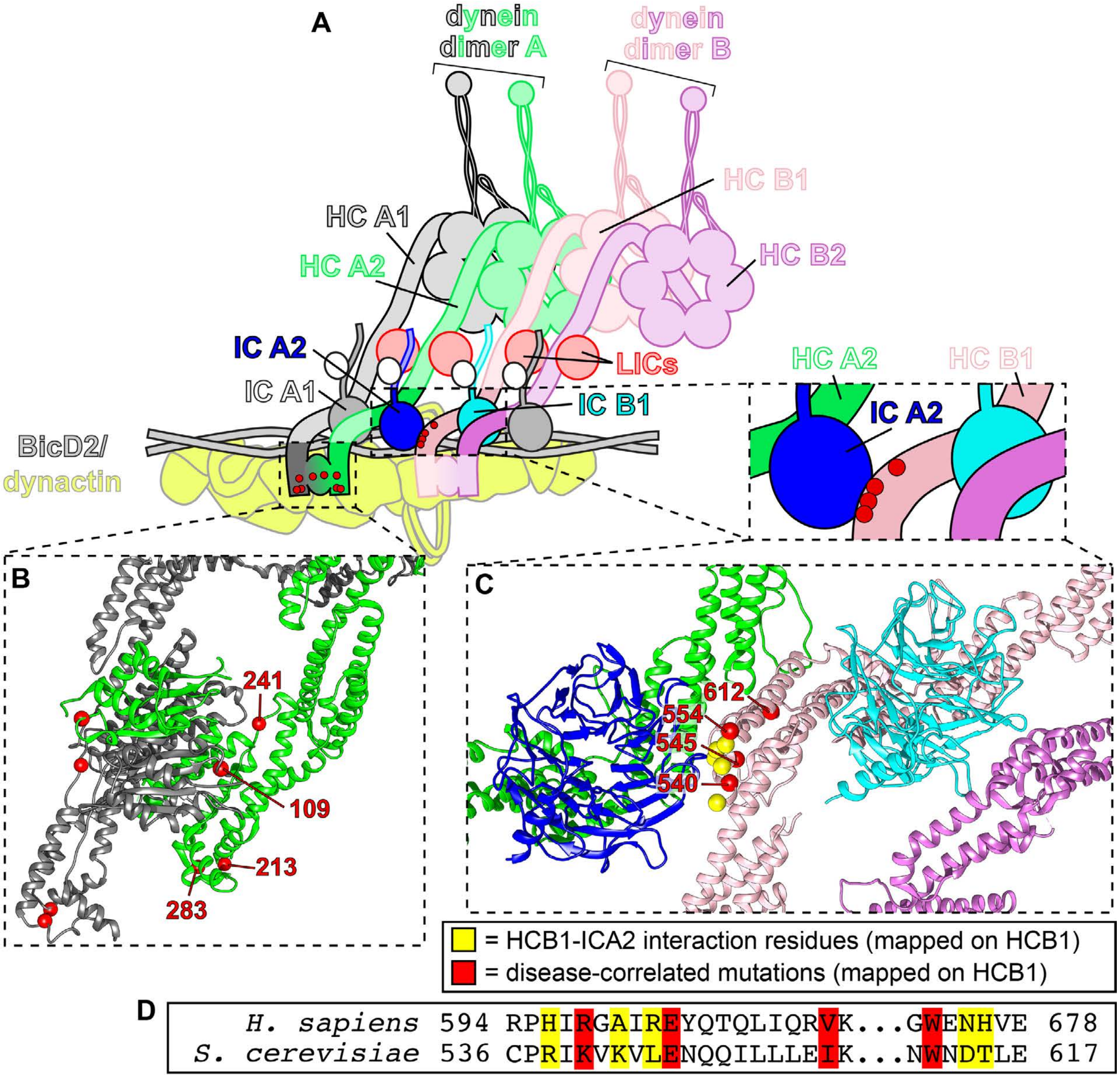

Figure 2, Supplement 4. **Tail domain mutations cluster to two distinct regions.** (A) Cartoon model depicting two dynein dimers (dimer A, composed of chains HCA1 and HCA2; and, dimer B, composed of chains HCB1 and HCB2) bound to a single dynactin complex with adapter BicDR1[3]. (B) Zoom-in view of the N-terminal dimerization domain of dynein (PDB 6F1T[3]) with red spheres denoting mutated residues (labeled on dynein heavy chain A2). (C) Zoom-in view of the contact point between dynein heavy chain HCB1 and the intermediate chain bound (ICA2) to heavy chain HCA2. Note that 3 of the 4 mutated residues cluster to the HCB1-ICA2 contact surface (K540, E545, I554), while one of them (W612) is somewhat distal from this region (red spheres, mutated residues; yellow spheres, contact points between HCB1 and ICA2). (D) Sequence alignment of the region shown in panel C.

#### Figure 3, Supplement 1

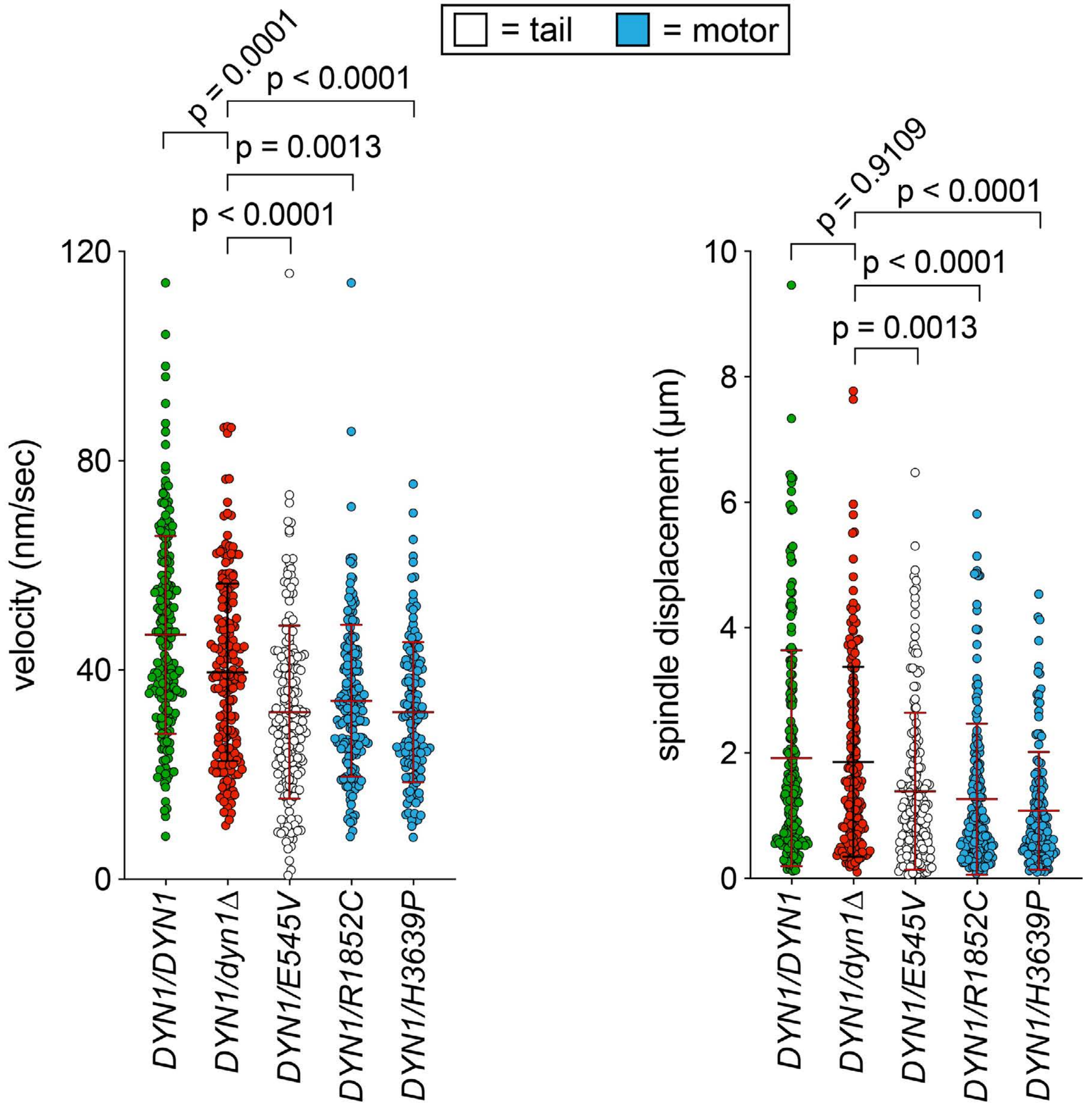

Figure 3, Supplement 1. **Additional plots of spindle dynamics data from diploid cells.** (A and B) Scatter plots depicting (A) velocity per spindle displacement event, and (B) displacement per spindle displacement event (from Figure 3). Mean values and standard deviations are also depicted with red and black lines.

### Figure 4, Supplement 1

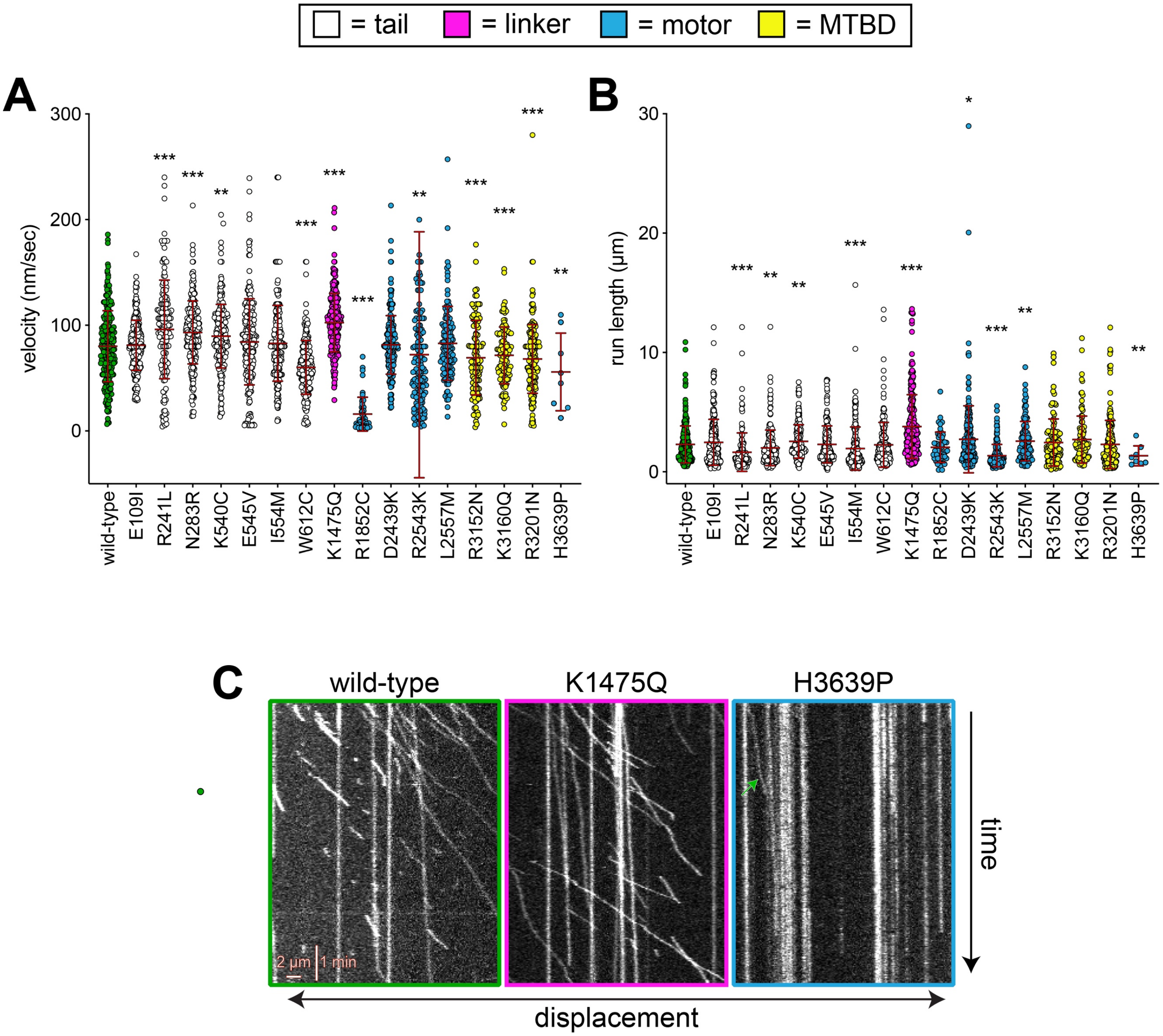

Figure 4, Supplement 1. **Additional plots of single molecule data and some representative kymographs.** (A and B) Scatter plots depicting (A) velocity, and (B) displacement values for single molecule motility assays (from Figure 4). Mean values and standard deviations are depicted with red lines. (C) Representative kymographs depicting single molecules of full-length dynein (wild-type or mutants, as indicated) walking along microtubules *in vitro*. Green arrow within H3639P panel depicts the only moving complex within the kymograph.

### Figure 4, Supplement 2

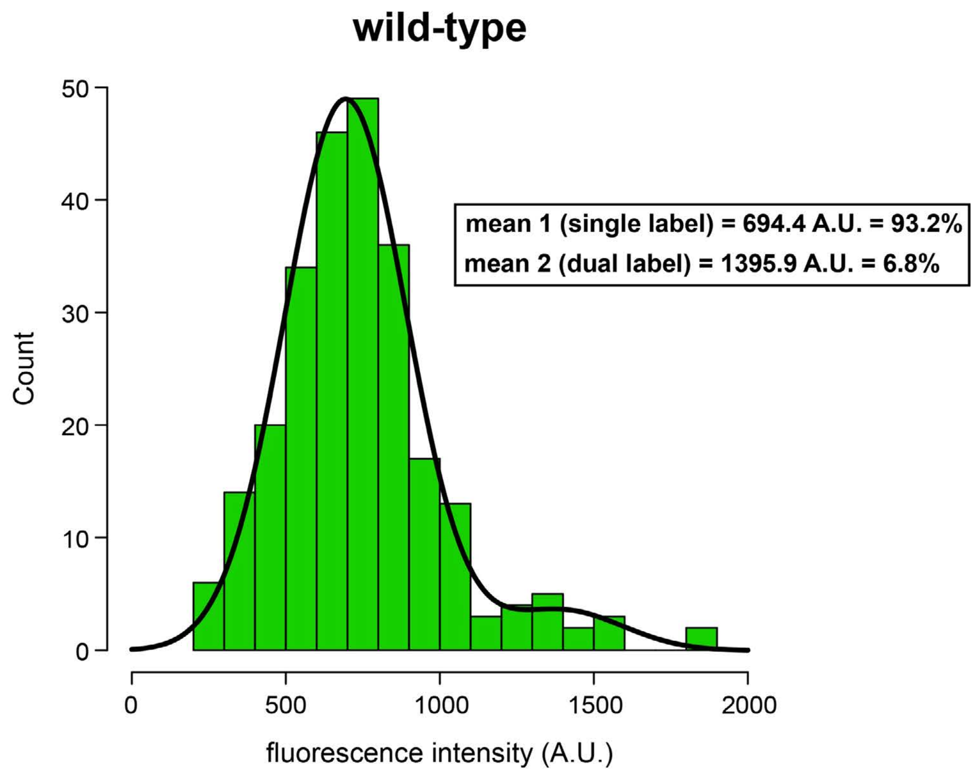


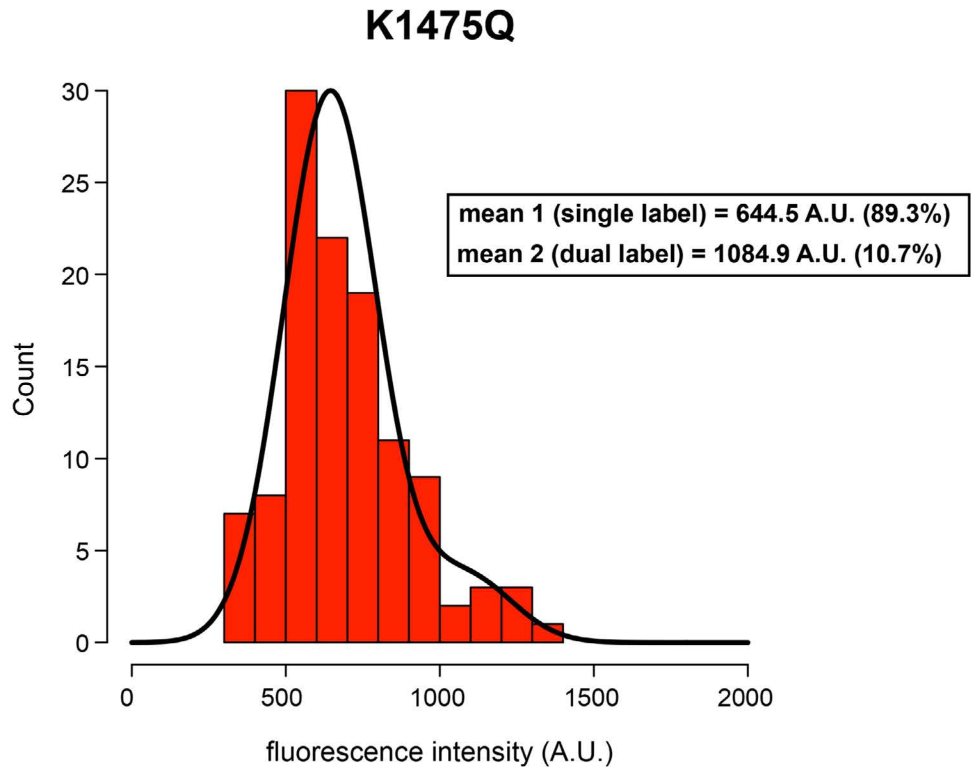

Figure 4, Supplement 2. **K1475Q mutant does not form aggregates in single molecule assay.** Histogram of fluorescence intensity values for single molecules of motile wild-type (top; n = 254 molecules) and K1475Q (bottom; n = 115 molecules) dynein, along with accompanying Gaussian fits and modeled parameters (determined using the model-based clustering algorithm Mclust[4]). The percentages reflect the relative proportion of molecules that fall within each component (*i.e.*, for mean 1, and mean 2). The two mean values for each likely represent single-labeled (mean 1) and dual-labeled (mean 2) dynein dimers, respectively. Importantly, the values for K1475Q are not higher than wild-type, indicating the increased processivity for this mutant is not a consequence of increased motor number.

### Figure 6, Supplement 1

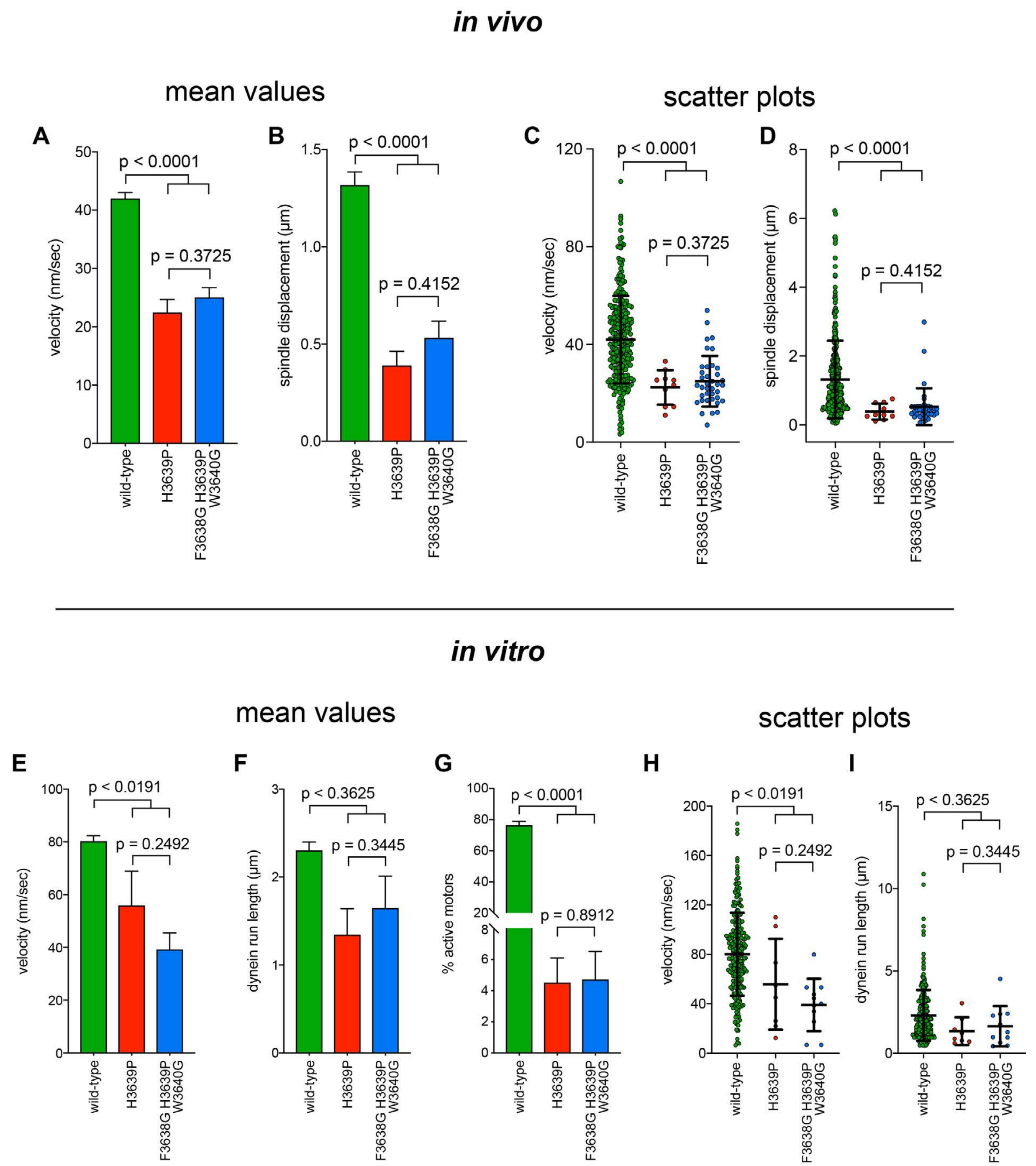

Figure 6, Supplement 1. **Additional insight into the molecular basis for dysfunction in H3639P.** (A - D) Plots depicting the velocity (A and C) and displacement (B and D; per event) values obtained from the spindle dynamics assay for the indicated haploid strains. For panels A and B, each data point represents the weighted mean ± weighted standard error (at least 29 HU-arrested cells from at least two independent experiments were analyzed for each strain). For panels C and D, mean values and standard deviations are depicted with black lines. (E - I) Quantitation of indicated parameters of single molecule motility. For panels E – G, each data point represents the weighted mean ± weighted standard error (E and F), or ± standard error of proportion (for G; at least 234 single molecules from at least two independent experiments were analyzed for each motor variant). For panels H and I, mean values and standard deviations are depicted with black lines. Statistical significance was determined using an unpaired Welch's t test (A, C, E and H), a Mann-Whitney test (B, D, F and I), or by calculating Z scores (G).

### Figure 7, Supplement 1

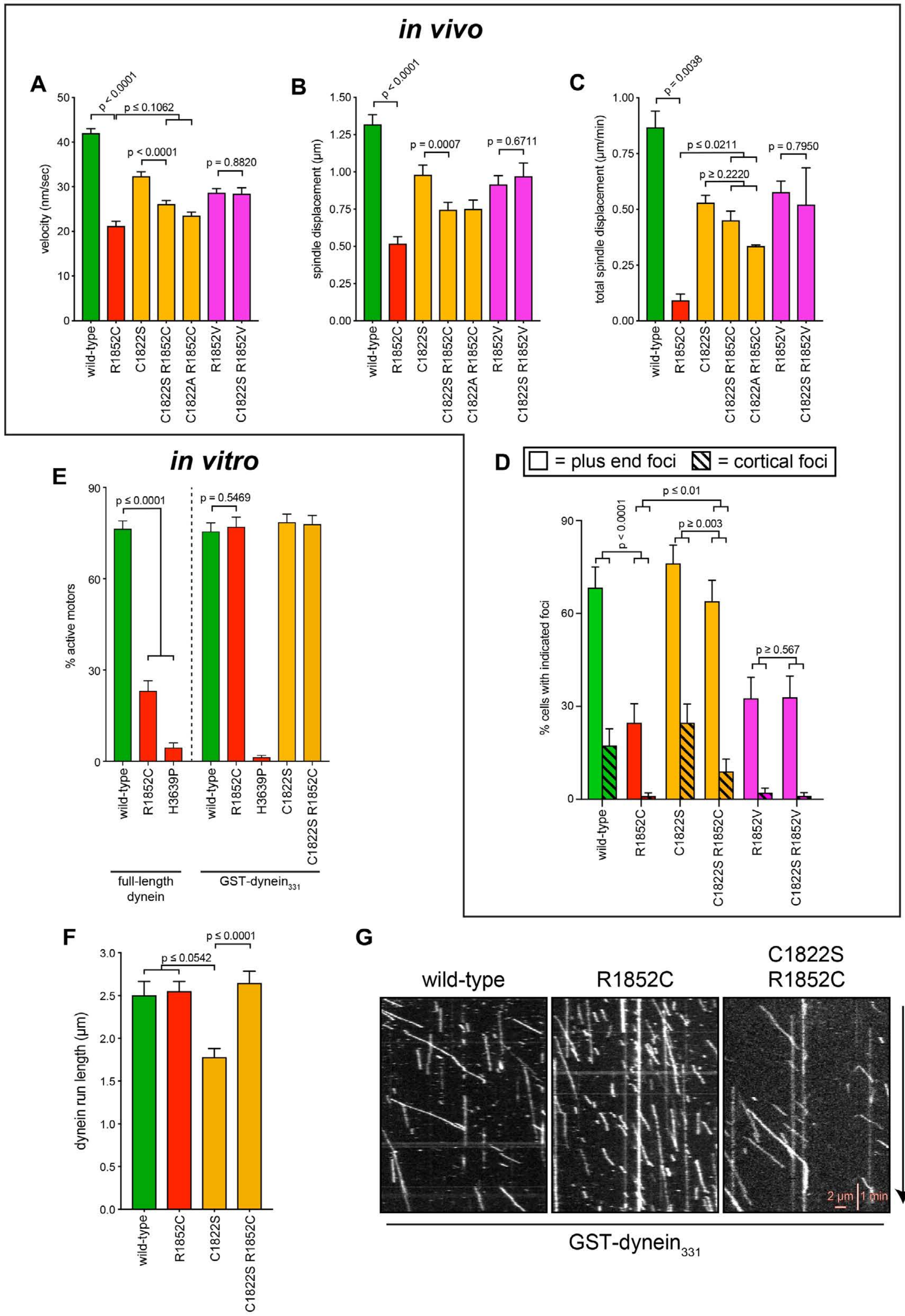

Figure 7, Supplement 1. **Additional insight into the molecular basis for dysfunction in R1852C.** (A - C) Plots depicting the indicated motility parameter values obtained from the spindle dynamics assay for the indicated haploid strains. Each data point represents the weighted mean ± weighted standard error (at least 28 HU-arrested cells from at least two independent experiments were analyzed for each strain). (D) The frequency of dynein localization to either microtubule plus ends or the cell cortex is plotted for indicated strains (scored as described in Figure 5 legend). Each data point represents the weighted mean ± weighted standard error (94 to 105 mitotic cells from at least two independent experiments were analyzed for each strain). (E) Plot depicting the fraction of active motors for the indicated full-length complexes (left), or the minimal motile fragment (GST-dynein$_{331}$; see text; right). Note the minimal fragment rescues the reduced fraction of active motors in the R1852C mutant. (F) Plot depicting single molecule run length values for indicated GST-dynein$_{331}$ variants. (G) Representative kymographs depicting single molecules of GST-dynein$_{331}$ (wild-type or mutants, as indicated) walking along microtubules *in vitro*. Statistical significance was determined using an unpaired Welch's t test (A, C), a Mann-Whitney test (B and F), or by calculating Z scores (D and E).

### Figure 7, Supplement 2

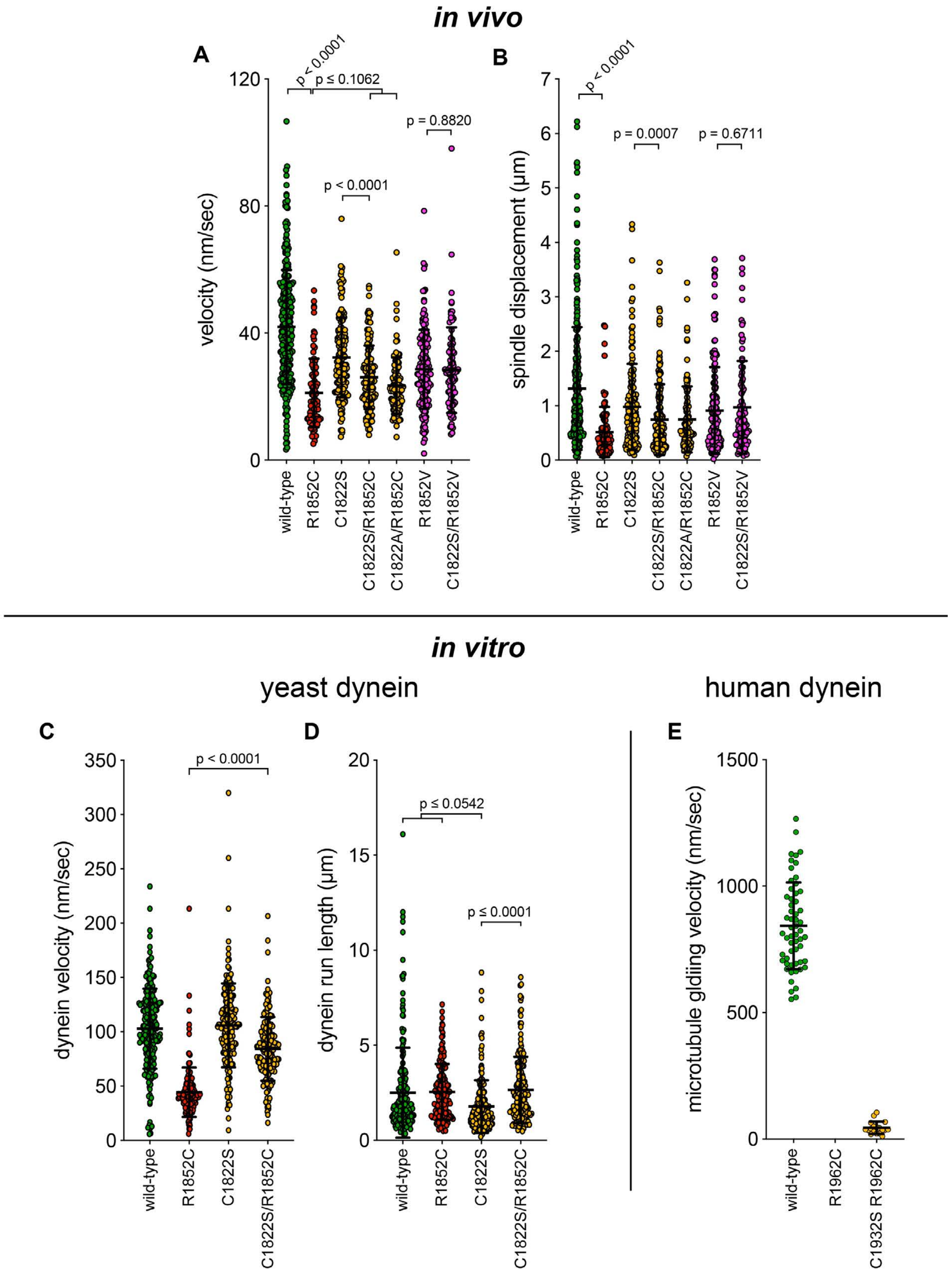

Figure 7, Supplement 2. **Additional plots of spindle dynamics and single molecule data.** (A and B) Scatter plots depicting the velocity and displacement (per event) obtained from the spindle dynamics assay for the indicated haploid strains (from Figure 7, Supplement 1). Mean values and standard deviations are depicted with black lines. (C - E) Scatter plots depicting velocity, and displacement values for single molecule motility assays from Figure 7 and Figure 7, Supplement 1. Mean values and standard deviations are depicted with black lines.

### Figure 8, Supplement 1

#### A. variance from wild-type

| mutation | | | | *in vivo* dynein-dynactin activity | | | | |
|---|---|---|---|---|---|---|---|---|
| | | | **Z score** | **q value** | **q value** | **q value** | **Z score** | **Z score** |
| human | yeast | disease | SP | V | D | ΣD | # events/min | NT |
| K129I | E109I | MCD | 1.180 | 0.062 | 2.128 | 0.122 | -2.416 | -0.319 |
| V236I | L213I | CMT | 0.451 | 7.591 | 1.007 | 0.445 | 2.299 | -2.577 |
| R264L | R241L | SMA-LED | 1.081 | 6.016 | 0.542 | 0.912 | -0.860 | -2.789 |
| H306R | N283R | SMA-LED | 1.158 | 4.566 | 0.322 | 0.189 | 1.280 | -1.238 |
| R598C | K540C | SMA-LED | ***-1.012*** | 0.990 | 1.965 | 2.079 | 1.995 | -0.100 |
| E603V | E545V | SMA-LED | 1.288 | 0.947 | 2.793 | 4.981 | -6.627 | -1.928 |
| V612M | I554M | SMA-LED | 1.011 | 2.993 | 1.068 | 0.812 | 0.143 | -0.659 |
| W673C | W612C | SMA-LED | 1.219 | 6.759 | 3.831 | 4.370 | -4.895 | -2.349 |
| R1567Q | K1475Q | MCD | 2.588 | 8.202 | 3.861 | 2.413 | -0.934 | -1.482 |
| R1962C | R1852C | MCD | 3.339 | 11.700 | 6.835 | 8.650 | -13.798 | -6.027 |
| E2616K | D2439K | SMA-LED | ***-0.991*** | 4.607 | 0.456 | 2.168 | -3.453 | -1.036 |
| R2720K | R2543K | MCD | 1.703 | 13.810 | 6.628 | 7.149 | -10.184 | -3.683 |
| V2734M | L2557M | CMD | 2.180 | 1.728 | 1.334 | 3.424 | -4.922 | -4.659 |
| K3336N | R3152N | MCD | 1.870 | 7.043 | 5.453 | 3.035 | 1.618 | -3.477 |
| R3344Q | K3160Q | MCD | 2.286 | 7.567 | 3.996 | 2.379 | 0.267 | -2.019 |
| R3384N | R3201N | MCD | 1.652 | 8.541 | 5.656 | 3.714 | -0.721 | -3.537 |
| H3822P | H3639P | MCD/SMA-LED | 12.276 | 4.085 | 2.949 | 9.540 | -19.195 | *i.o.* |
| | *dyn1Δ* | | 11.900 | 33.960 | 16.180 | 9.629 | -20.584 | - |

#### B. normalized relative variance (nrv) from wild-type

| mutation | | | *in vivo* dynein-dynactin activity | | | | | |
|---|---|---|---|---|---|---|---|---|
| human | yeast | disease | SP | V | D | ΣD | # events/min | NT |
| K129I | E109I | MCD | 0.481 | 0.002 | 0.132 | 0.013 | 0.117 | 0.053 |
| V236I | L213I | CMT | 0.184 | 0.224 | 0.062 | 0.046 | 0.112 | 0.428 |
| R264L | R241L | SMA-LED | 0.440 | 0.177 | 0.033 | 0.095 | 0.042 | 0.463 |
| H306R | N283R | SMA-LED | 0.471 | 0.134 | 0.020 | 0.020 | 0.062 | 0.205 |
| R598C | K540C | SMA-LED | ***0.000*** | 0.029 | 0.121 | 0.216 | 0.097 | 0.017 |
| E603V | E545V | SMA-LED | 0.524 | 0.028 | 0.173 | 0.517 | 0.322 | 0.320 |
| V612M | I554M | SMA-LED | 0.412 | 0.088 | 0.066 | 0.084 | 0.007 | 0.109 |
| W673C | W612C | SMA-LED | 0.496 | 0.199 | 0.237 | 0.454 | 0.238 | 0.390 |
| R1567Q | K1475Q | MCD | 1.054 | 0.242 | 0.239 | 0.251 | 0.045 | 0.246 |
| R1962C | R1852C | MCD | 1.360 | 0.345 | 0.422 | 0.898 | 0.670 | 1.000 |
| E2616K | D2439K | SMA-LED | ***0.000*** | 0.136 | 0.028 | 0.225 | 0.168 | 0.172 |
| R2720K | R2543K | MCD | 0.694 | 0.407 | 0.410 | 0.742 | 0.495 | 0.611 |
| V2734M | L2557M | CMD | 0.888 | 0.051 | 0.082 | 0.356 | 0.239 | 0.773 |
| K3336N | R3152N | MCD | 0.762 | 0.207 | 0.337 | 0.315 | 0.079 | 0.577 |
| R3344Q | K3160Q | MCD | 0.931 | 0.223 | 0.247 | 0.247 | 0.013 | 0.335 |
| R3384N | R3201N | MCD | 0.673 | 0.252 | 0.350 | 0.386 | 0.035 | 0.587 |
| H3822P | H3639P | MCD/SMA-LED | 5.000 | 0.120 | 0.182 | 0.991 | 0.933 | - |
| | *dyn1Δ* | | 4.847 | 1.000 | 1.000 | 1.000 | 1.000 | - |
| | wild-type | | 0.000 | 0.000 | 0.000 | 0.000 | 0.000 | 0.000 |

#### C. CDD calculation

$$nrv = \frac{|v|}{v_{max}}$$

$$CDD = \frac{(5 \cdot nrv_{SP}) + nrv_{V} + nrv_{D} + nrv_{\Sigma D} + nrv_{\#/min} + nrv_{NT}}{6}$$

Figure 8, Supplement 1. **Data used for tabulation of the coefficient of dynein dysfunction (CDD).** (A) Z scores (see methods) and q values for differences between mutant and wild-type cells for each assay. Graphpad Prism was used to calculate q values (the difference between the two means divided by the standard error of that difference). (B) Relative difference between mean values (Z scores and q values, "v") for mutant and wild-type cells are expressed as normalized relative variance (nrv), where nrv = $|v|/v_{max}$ for each column. Colors indicate relative degree of difference between mutant and wild-type for each value. (C) Coefficient of dynein dysfunction (CDD) was calculated from the values shown in panel B. In the two cases where a value wasn't determined (due to insufficient observations), the denominator was reduced from 6 to 5. Note that in two instances, the Z score for spindle positioning was negative (due to a lower number of mispositioned spindles being observed in K540C and D2439K cells than in wild-type cells; see Fig. 1B). These two values were corrected to 0 so as to avoid them skewing the $nrv_{SP}$ values. Also see Methods.

**TABLE S1.** Strains used throughout this study.

| Strain number | Genotype | Yeast background | Source |
|---|---|---|---|
| SMY1008 | Mata *GAL1p:ZZ-TEV-6HIS-GFP-3HA-GST-dyn1$_{331}$-HaloTag::KAN$^R$ prb1Δ pep4Δ::HIS5 his3-11,15 ura3-52 leu2-3,112 ade2-1 trp-1* | W303 | Ref[5] |
| SMY1010 | Mata *DYN1-3GFP::TRP1 TUB1+3'UTR::HPH::HIS3p:mRuby2-TUB1 ura3-52 lys2-801 leu2-Δ1 his3-Δ200 trp1-Δ63* | YEF473 | This study |
| SMY1066 | Mata *dyn1$^{E109I}$-3GFP::TRP1 TUB1+3'UTR::HPH::HIS3p:mRuby2-TUB1 ura3-52 lys2-801 leu2-Δ1 his3-Δ200 trp1-Δ63* | YEF473 | This study |
| SMY1068 | Mata *dyn1$^{L213I}$-3GFP::TRP1 TUB1+3'UTR::HPH::HIS3p:mRuby2-TUB1 ura3-52 lys2-801 leu2-Δ1 his3-Δ200 trp1-Δ63* | YEF473 | This study |

| | | | |
|---|---|---|---|
| SMY1073 | Mata *dyn1$^{W612C}$-3GFP::TRP1 TUB1+3'UTR::HPH::HIS3p:mRuby2-TUB1 ura3-52 lys2-801 leu2-Δ1 his3-Δ200 trp1-Δ63* | YEF473 | This study |
| SMY1074 | Mata *dyn1$^{W612C}$-3GFP::TRP1 TUB1+3'UTR::HPH::HIS3p:mRuby2-TUB1 ura3-52 lys2-801 leu2-Δ1 his3-Δ200 trp1-Δ63* | YEF473 | This study |
| SMY1089 | Mata *dyn1$^{K540C}$-3GFP::TRP1 TUB1+3'UTR::HPH::HIS3p:mRuby2-TUB1 ura3-52 lys2-801 leu2-Δ1 his3-Δ200 trp1-Δ63* | YEF473 | This study |
| SMY1090 | Mata *dyn1$^{K3160Q}$-3GFP::TRP1 TUB1+3'UTR::HPH::HIS3p:mRuby2-TUB1 ura3-52 lys2-801 leu2-Δ1 his3-Δ200 trp1-Δ63* | YEF473 | This study |
| SMY1105 | Mata *dyn1$^{D2439K}$-3GFP::TRP1 TUB1+3'UTR::HPH::HIS3p:mRuby2-TUB1 ura3-52 lys2-801 leu2-Δ1 his3-Δ200 trp1-Δ63* | YEF473 | This study |
| SMY1107 | Mata *dyn1$^{H3639P}$-3GFP::TRP1 TUB1+3'UTR::HPH::HIS3p:mRuby2-TUB1 ura3-52 lys2-801 leu2-Δ1 his3-Δ200 trp1-Δ63* | YEF473 | This study |
| SMY1111 | Mata *dyn1$^{E545V}$-3GFP::TRP1 TUB1+3'UTR::HPH::HIS3p:mRuby2-TUB1 ura3-52 lys2-801 leu2-Δ1 his3-Δ200 trp1-Δ63* | YEF473 | This study |
| SMY1112 | Mata *dyn1$^{E545V}$-3GFP::TRP1 TUB1+3'UTR::HPH::HIS3p:mRuby2-TUB1 ura3-52 lys2-801 leu2-Δ1 his3-Δ200 trp1-Δ63* | YEF473 | This study |
| SMY1113 | Mata *dyn1$^{I554M}$-3GFP::TRP1 TUB1+3'UTR::HPH::HIS3p:mRuby2-TUB1 ura3-52 lys2-801 leu2-Δ1 his3-Δ200 trp1-Δ63* | YEF473 | This study |
| SMY1116 | Mata *dyn1$^{K1475Q}$-3GFP::TRP1 TUB1+3'UTR::HPH::HIS3p:mRuby2-TUB1 ura3-52 lys2-801 leu2-Δ1 his3-Δ200 trp1-Δ63* | YEF473 | This study |
| SMY1117 | Mata *dyn1$^{L2557M}$-3GFP::TRP1 TUB1+3'UTR::HPH::HIS3p:mRuby2-TUB1 ura3-52 lys2-801 leu2-Δ1 his3-Δ200 trp1-Δ63* | YEF473 | This study |
| SMY1119 | Mata *dyn1$^{R3152N}$-3GFP::TRP1 TUB1+3'UTR::HPH::HIS3p:mRuby2-TUB1 ura3-52 lys2-801 leu2-Δ1 his3-Δ200 trp1-Δ63* | YEF473 | This study |
| SMY1160 | Matα *DYN1-3GFP::TRP1 kar9Δ::KAN$^{R}$ GFP-TUB1::LEU2 ura3-52 lys2- 801 leu2-Δ1 his3-Δ200 trp1-Δ63* | YEF473 | This study |
| SMY1161 | Mata *Dyn1-3GFP::TRP1 kar9Δ::KAN$^{R}$ GFP-TUB1::LEU2 ura3-52 lys2- 801 leu2-Δ1 his3-Δ200 trp1-Δ63* | YEF473 | This study |

| | | | |
|---|---|---|---|
| SMY1178 | Mata *dyn1$^{R3201N}$-3GFP::TRP1 TUB1+3'UTR::HPH::HIS3p:mRuby2-TUB1 ura3-52 lys2-801 leu2-Δ1 his3-Δ200 trp1-Δ63* | YEF473 | This study |
| SMY1179 | Mata *dyn1$^{R3201N}$-3GFP::TRP1 TUB1+3'UTR::HPH::HIS3p:mRuby2-TUB1 ura3-52 lys2-801 leu2-Δ1 his3-Δ200 trp1-Δ63* | YEF473 | This study |
| SMY1185 | Mata *Dyn1-3GFP::TRP1 NUP133-3mCherry::URA3 kar9Δ::KAN$^{R}$ GFP-TUB1::LEU2 ura3-52 lys2- 801 leu2-Δ1 his3-Δ200 trp1-Δ63* | YEF473 | This study |
| SMY1186 | *Mata Dyn1-3GFP::TRP1 NUP133-3mCherry::URA3 kar9Δ::KAN$^{R}$ GFP-TUB1::LEU2 ura3-52 lys2- 801 leu2-Δ1 his3-Δ200 trp1-Δ63* | YEF473 | This study |
| SMY1196 | *Matα dyn1$^{E545V}$-3GFP::TRP1 NUP133-3mCherry::URA3 kar9Δ::KAN$^{R}$ GFP-TUB1::LEU2 ura3-52 lys2- 801 leu2-Δ1 his3-Δ200 trp1-Δ63* | YEF473 | This study |
| SMY1198 | *Mata dyn1$^{E109I}$-3GFP::TRP1 NUP133-3mCherry::URA3 kar9Δ::KAN$^{R}$ GFP-TUB1::LEU2 ura3-52 lys2- 801 leu2-Δ1 his3-Δ200 trp1-Δ63* | YEF473 | This study |
| SMY1200 | Matα *dyn1$^{L2557M}$-3GFP::TRP1 NUP133-3mCherry::URA3 kar9Δ::KAN$^{R}$ GFP-TUB1::LEU2 ura3-52 lys2- 801 leu2-Δ1 his3-Δ200 trp1-Δ63* | YEF473 | This study |
| SMY1201 | Matα *dyn1$^{L2557M}$-3GFP::TRP1 NUP133-3mCherry::URA3 kar9Δ::KAN$^{R}$ GFP-TUB1::LEU2 ura3-52 lys2- 801 leu2-Δ1 his3-Δ200 trp1-Δ63* | YEF473 | This study |
| SMY1220 | Mata *dyn1$^{R2439K}$-3GFP::TRP1 NUP133-3mCherry::URA3 kar9Δ::KAN$^{R}$ GFP-TUB1::LEU2 ura3-52 lys2- 801 leu2-Δ1 his3-Δ200 trp1-Δ63* | YEF473 | This study |
| SMY1222 | Mata *dyn1$^{K540C}$-3GFP::TRP1 NUP133-3mCherry::URA3 kar9Δ::KAN$^{R}$ GFP-TUB1::LEU2 ura3-52 lys2- 801 leu2-Δ1 his3-Δ200 trp1-Δ63* | YEF473 | This study |
| SMY1243 | Mata *dyn1$^{H3639P}$-3GFP::TRP1 TUB1+3'UTR::HPH::HIS3p:mRuby2-TUB1 ura3-52 lys2-801 leu2-Δ1 his3-Δ200 trp1-Δ63* | YEF473 | This study |
| SMY1254 | Matα *dyn1$^{R3152N}$-3GFP::TRP1 NUP133-3mCherry::URA3 kar9Δ::KAN$^{R}$ GFP-TUB1::LEU2 ura3-52 lys2- 801 leu2-Δ1 his3-Δ200 trp1-Δ63* | YEF473 | This study |
| SMY1266 | Mata *DYN3-13MYC::HPH PAC11-13MYC::TRP ZZ-TEV-3HA-DYN1-HaloTag::KAN$^{R}$ nip100Δ pep4Δ::HIS5 prb1Δ his3-11,15 ura3-52 leu2-3,112 ade2-1 trp-1* | W303 | This study |
| SMY1317 | Mata *dyn1$^{R1852C}$-3GFP::TRP1 TUB1+3'UTR::HPH::HIS3p:mRuby2-TUB1 ura3-52 lys2-801 leu2-Δ1 his3-Δ200 trp1-Δ63* | YEF473 | This study |

| | | | |
|---|---|---|---|
| SMY1318 | Mata *dyn1*$^{R3201N}$*-3GFP::TRP1 TUB1+3'UTR::HPH::HIS3p:mRuby2-TUB1 ura3-52 lys2-801 leu2-Δ1 his3-Δ200 trp1-Δ63* | YEF473 | This study |
| SMY1327 | Mata *dyn1*$^{R241L}$*-3GFP::TRP1 TUB1+3'UTR::HPH::HIS3p:mRuby2-TUB1 ura3-52 lys2-801 leu2-Δ1 his3-Δ200 trp1-Δ63* | YEF473 | This study |
| SMY1351 | Mata *dyn1*$^{R1852C}$*-3GFP::TRP1 NUP133-3mCherry::URA3 kar9Δ::KAN*$^{R}$ *GFP-TUB1::LEU2 ura3-52 lys2- 801 leu2-Δ1 his3-Δ200 trp1-Δ63* | YEF473 | This study |
| SMY1369 | Mata *dyn1*$^{W3640P}$*-3GFP::TRP1 TUB1+3'UTR::HPH::HIS3p:mRuby2-TUB1 ura3-52 lys2-801 leu2-Δ1 his3-Δ200 trp1-Δ63* | YEF473 | This study |
| SMY1370 | Mata *dyn1*$^{W3640P}$*-3GFP::TRP1 TUB1+3'UTR::HPH::HIS3p:mRuby2-TUB1 ura3-52 lys2-801 leu2-Δ1 his3-Δ200 trp1-Δ63* | YEF473 | This study |
| SMY1371 | Mata *dyn1*$^{F3641P}$*-3GFP::TRP1 TUB1+3'UTR::HPH::HIS3p:mRuby2-TUB1 ura3-52 lys2-801 leu2-Δ1 his3-Δ200 trp1-Δ63* | YEF473 | This study |
| SMY1372 | Mata *dyn1*$^{F3641P}$*-3GFP::TRP1 TUB1+3'UTR::HPH::HIS3p:mRuby2-TUB1 ura3-52 lys2-801 leu2-Δ1 his3-Δ200 trp1-Δ63* | YEF473 | This study |
| SMY1373 | Mata *dyn1*$^{Y3642P}$*-3GFP::TRP1 TUB1+3'UTR::HPH::HIS3p:mRuby2-TUB1 ura3-52 lys2-801 leu2-Δ1 his3-Δ200 trp1-Δ63* | YEF473 | This study |
| SMY1374 | Mata *dyn1*$^{I3644P}$*-3GFP::TRP1 TUB1+3'UTR::HPH::HIS3p:mRuby2-TUB1 ura3-52 lys2-801 leu2-Δ1 his3-Δ200 trp1-Δ63* | YEF473 | This study |
| SMY1381 | Mata *dyn1*$^{N283R}$*-3GFP::TRP1 TUB1+3'UTR::HPH::HIS3p:mRuby2-TUB1 ura3-52 lys2-801 leu2-Δ1 his3-Δ200 trp1-Δ63* | YEF473 | This study |
| SMY1386 | Mata *dyn1*$^{R3201N}$*-3GFP::TRP1 NUP133-3mCherry::URA3 kar9Δ::KAN*$^{R}$ *GFP-TUB1::LEU2 ura3-52 lys2- 801 leu2-Δ1 his3-Δ200 trp1-Δ63* | YEF473 | This study |
| SMY1420 | Mata *dyn1*$^{G3643P}$*-3GFP::TRP1 TUB1+3'UTR::HPH::HIS3p:mRuby2-TUB1 ura3-52 lys2-801 leu2-Δ1 his3-Δ200 trp1-Δ63* | YEF473 | This study |
| SMY1443 | Mata *DYN3-13MYC::HPH PAC11-13MYC::TRP ZZ-TEV-3HA-dyn1*$^{R3201N}$*-HaloTag::KAN*$^{R}$ *nip100Δ pep4Δ::HIS5 prb1Δ his3-11,15 ura3-52 leu2-3,112 ade2-1 trp-1* | W303 | This study |
| SMY1444 | Mata *DYN3-13MYC::HPH PAC11-13MYC::TRP ZZ-TEV-3HA-dyn1*$^{R3201N}$*-HaloTag::KAN*$^{R}$ *nip100Δ pep4Δ::HIS5 prb1Δ his3-11,15 ura3-52 leu2-3,112 ade2-1 trp-1* | W303 | This study |

| | | | |
|---|---|---|---|
| SMY1445 | Mata *DYN3-13MYC::HPH PAC11-13MYC::TRP ZZ-TEV-3HA-dyn1*$^{D2439K}$*-HaloTag::KAN*$^{R}$ *nip100Δ pep4Δ::HIS5 prb1Δ his3-11,15 ura3-52 leu2-3,112 ade2-1 trp-1* | W303 | This study |
| SMY1447 | Mata *DYN3-13MYC::HPH PAC11-13MYC::TRP ZZ-TEV-3HA-dyn1*$^{L2557M}$*-HaloTag::KAN*$^{R}$ *nip100Δ pep4Δ::HIS5 prb1Δ his3-11,15 ura3-52 leu2-3,112 ade2-1 trp-1* | W303 | This study |
| SMY1448 | Mata *DYN3-13MYC::HPH PAC11-13MYC::TRP ZZ-TEV-3HA-dyn1*$^{K1475Q}$*-HaloTag::KAN*$^{R}$ *nip100Δ pep4Δ::HIS5 prb1Δ his3-11,15 ura3-52 leu2-3,112 ade2-1 trp-1* | W303 | This study |
| SMY1449 | Mata *DYN3-13MYC::HPH PAC11-13MYC::TRP ZZ-TEV-3HA-dyn1*$^{W612C}$*-HaloTag::KAN*$^{R}$ *nip100Δ pep4Δ::HIS5 prb1Δ his3-11,15 ura3-52 leu2-3,112 ade2-1 trp-1* | W303 | This study |
| SMY1455 | Mata *DYN3-13MYC::HPH PAC11-13MYC::TRP ZZ-TEV-3HA-dyn1*$^{H3639P}$*-HaloTag::KAN*$^{R}$ *nip100Δ pep4Δ::HIS5 prb1Δ his3-11,15 ura3-52 leu2-3,112 ade2-1 trp-1* | W303 | This study |
| SMY1456 | Mata *DYN3-13MYC::HPH PAC11-13MYC::TRP ZZ-TEV-3HA-dyn1*$^{E545V}$*-HaloTag::KAN*$^{R}$ *nip100Δ pep4Δ::HIS5 prb1Δ his3-11,15 ura3-52 leu2-3,112 ade2-1 trp-1* | W303 | This study |
| SMY1458 | Matα/Mata *DYN1-3GFP::TRP1/ DYN1-3GFP::TRP1 NUP133-3mCherry::URA3/NUP133-3mCherry::URA3 kar9Δ::KAN*$^{R}$*/kar9Δ::KAN*$^{R}$ *GFP-TUB1::LEU2/GFP-TUB1::LEU2 ura3-52/ura3-52 lys2- 801/lys2- 801 leu2-Δ1/leu2-Δ1 his3-Δ200/his3-Δ200 trp1-Δ63/trp1-Δ63* | YEF473 | This study |
| SMY1481 | Mata *DYN3-13MYC::HPH PAC11-13MYC::TRP ZZ-TEV-3HA-dyn1*$^{R241L}$*-HaloTag::KAN*$^{R}$ *nip100Δ pep4Δ::HIS5 prb1Δ his3-11,15 ura3-52 leu2-3,112 ade2-1 trp-1* | W303 | This study |
| SMY1507 | Mata *dyn1*$^{C1822S}$*-3GFP::TRP1 TUB1+3'UTR::HPH::HIS3p:mRuby2-TUB1 ura3-52 lys2-801 leu2-Δ1 his3-Δ200 trp1-Δ63* | YEF473 | This study |
| SMY1508 | Mata *dyn1*$^{R1822S,R1852C}$*-3GFP::TRP1 TUB1+3'UTR::HPH::HIS3p:mRuby2-TUB1 ura3-52 lys2-801 leu2-Δ1 his3-Δ200 trp1-Δ63* | YEF473 | This study |
| SMY1509 | Mata *dyn1*$^{R1822S,R1852C}$*-3GFP::TRP1 TUB1+3'UTR::HPH::HIS3p:mRuby2-TUB1 ura3-52 lys2-801 leu2-Δ1 his3-Δ200 trp1-Δ63* | YEF473 | This study |

| | | | |
|---|---|---|---|
| SMY1520 | Mata *DYN3-13MYC::HPH PAC11-13MYC::TRP ZZ-TEV-3HA-dyn1*$^{K3160Q}$*-HaloTag::KAN*$^{R}$ *nip100Δ pep4Δ::HIS5 prb1Δ his3-11,15 ura3-52 leu2-3,112 ade2-1 trp-1* | W303 | This study |
| SMY1532 | Mata *dyn1*$^{C1822S,R1852C}$*-3GFP::TRP1 NUP133-3mCherry::URA3 kar9Δ::KAN*$^{R}$ *GFP-TUB1::LEU2 ura3-52 lys2- 801 leu2-Δ1 his3-Δ200 trp1-Δ63* | YEF473 | This study |
| SMY1533 | Mata *dyn1*$^{N283R}$*-3GFP::TRP1 NUP133-3mCherry::URA3 kar9Δ::KAN*$^{R}$ *GFP-TUB1::LEU2 ura3-52 lys2- 801 leu2-Δ1 his3-Δ200 trp1-Δ63* | YEF473 | This study |
| SMY1545 | Mata *DYN3-13MYC::HPH PAC11-13MYC::TRP ZZ-TEV-3HA-dyn1*$^{K540C}$*-HaloTag::KAN*$^{R}$ *nip100Δ pep4Δ::HIS5 prb1Δ his3-11,15 ura3-52 leu2-3,112 ade2-1 trp-1* | W303 | This study |
| SMY1546 | Mata *DYN3-13MYC::HPH PAC11-13MYC::TRP ZZ-TEV-3HA-dyn1*$^{K540C}$*-HaloTag::KAN*$^{R}$ *nip100Δ pep4Δ::HIS5 prb1Δ his3-11,15 ura3-52 leu2-3,112 ade2-1 trp-1* | W303 | This study |
| SMY1547 | Mata *DYN3-13MYC::HPH PAC11-13MYC::TRP ZZ-TEV-3HA-dyn1*$^{E545V}$*-HaloTag::KAN*$^{R}$ *nip100Δ pep4Δ::HIS5 prb1Δ his3-11,15 ura3-52 leu2-3,112 ade2-1 trp-1* | W303 | This study |
| SMY1565 | Matα *dyn1*$^{C1822S,R1852C}$*-3GFP::TRP1 NUP133-3mCherry::URA3 kar9Δ::KAN*$^{R}$ *GFP-TUB1::LEU2 ura3-52 lys2- 801 leu2-Δ1 his3-Δ200 trp1-Δ63* | YEF473 | This study |
| SMY1585 | Mata *GAL1p:ZZ-TEV-6HIS-GFP-3HA-GST-dyn1*$_{331}^{C1822S}$*-HaloTag::KAN*$^{R}$ *prb1Δ pep4Δ::HIS5 his3-11,15 ura3-52 leu2-3,112 ade2-1 trp-1* | W303 | This study |
| SMY1588 | Mata *GAL1p:ZZ-TEV-6HIS-GFP-3HA-GST-dyn1*$_{331}^{R1852C}$*-HaloTag::KAN*$^{R}$ *prb1Δ pep4Δ::HIS5 his3-11,15 ura3-52 leu2-3,112 ade2-1 trp-1* | W303 | This study |
| SMY1589 | Mata *GAL1p:ZZ-TEV-6HIS-GFP-3HA-GST-dyn1*$_{331}^{R1852C}$*-HaloTag::KAN*$^{R}$ *prb1Δ pep4Δ::HIS5 his3-11,15 ura3-52 leu2-3,112 ade2-1 trp-1* | W303 | This study |
| SMY1591 | Matα *dyn1*$^{H3639P}$*-3GFP::TRP1 NUP133-3mCherry::URA3 kar9Δ::KAN*$^{R}$ *GFP-TUB1::LEU2 ura3-52 lys2- 801 leu2-Δ1 his3-Δ200 trp1-Δ63* | YEF473 | This study |
| SMY1592 | Matα/Mata *DYN1-3GFP::TRP1/ dyn1Δ::HIS3 NUP133-3mCherry::URA3/NUP133-3mCherry::URA3 kar9Δ::KAN*$^{R}$*/kar9Δ::KAN*$^{R}$ *GFP-TUB1::LEU2/GFP-TUB1::LEU2 ura3-52/ura3-52 lys2- 801/lys2- 801 leu2-Δ1/leu2-Δ1 his3-Δ200/his3-Δ200 trp1-Δ63/trp1-Δ63* | YEF473 | This study |

| | | | |
|---|---|---|---|
| SMY1627 | Mata *dyn1*$^{R2543K}$*-3GFP::TRP1 TUB1+3'UTR::HPH::HIS3p:mRuby2-TUB1 ura3-52 lys2-801 leu2-Δ1 his3-Δ200 trp1-Δ63* | YEF473 | This study |
| SMY1628 | Mata *dyn1*$^{R2543K}$*-3GFP::TRP1 TUB1+3'UTR::HPH::HIS3p:mRuby2-TUB1 ura3-52 lys2-801 leu2-Δ1 his3-Δ200 trp1-Δ63* | YEF473 | This study |
| SMY1651 | Mata *GAL1p:ZZ-TEV-6HIS-GFP-3HA-GST-dyn1*$_{331}^{H3639P}$*-HaloTag::KAN*$^{R}$ *prb1Δ pep4Δ::HIS5 his3-11,15 ura3-52 leu2-3,112 ade2-1 trp-1* | W303 | This study |
| SMY1678 | Matα/Mata *DYN1/ dyn1*$^{E545V}$*::TRP1 NUP133-3mCherry::URA3/NUP133-3mCherry::URA3 kar9Δ::KAN*$^{R}$*/kar9Δ::KAN*$^{R}$ *GFP-TUB1::LEU2/GFP-TUB1::LEU2 ura3-52/ura3-52 lys2- 801/lys2- 801 leu2-Δ1/leu2-Δ1 his3-Δ200/his3-Δ200 trp1-Δ63/trp1-Δ63* | YEF473 | This study |
| SMY1679 | Matα/Mata *DYN1/ dyn1*$^{H3639P}$*::TRP1 NUP133-3mCherry::URA3/NUP133-3mCherry::URA3 kar9Δ::KAN*$^{R}$*/kar9Δ::KAN*$^{R}$ *GFP-TUB1::LEU2/GFP-TUB1::LEU2 ura3-52/ura3-52 lys2- 801/lys2- 801 leu2-Δ1/leu2-Δ1 his3-Δ200/his3-Δ200 trp1-Δ63/trp1-Δ63* | YEF473 | This study |
| SMY1697 | Mata *dyn1*$^{C1822S}$*-3GFP::TRP1 NUP133-3mCherry::URA3 kar9Δ::KAN*$^{R}$ *GFP-TUB1::LEU2 ura3-52 lys2- 801 leu2-Δ1 his3-Δ200 trp1-Δ63* | YEF473 | This study |
| SMY1698 | Mata *GAL1p:ZZ-TEV-6HIS-GFP-3HA-GST-dyn1*$_{331}^{C1822S,R1852C}$*-HaloTag::KAN*$^{R}$ *prb1Δ pep4Δ::HIS5 his3-11,15 ura3-52 leu2-3,112 ade2-1 trp-1* | W303 | This study |
| SMY1699 | Mata *GAL1p:ZZ-TEV-6HIS-GFP-3HA-GST-dyn1*$_{331}^{C1822S,R1852C}$*-HaloTag::KAN*$^{R}$ *prb1Δ pep4Δ::HIS5 his3-11,15 ura3-52 leu2-3,112 ade2-1 trp-1* | W303 | This study |
| SMY1727 | Matα *dyn1*$^{I554M}$*-3GFP::TRP1 NUP133-3mCherry::URA3 kar9Δ::KAN*$^{R}$ *GFP-TUB1::LEU2 ura3-52 lys2- 801 leu2-Δ1 his3-Δ200 trp1-Δ63* | YEF473 | This study |
| SMY1732 | Mata *dyn1*$^{R2543K}$*-3GFP::TRP1 NUP133-3mCherry::URA3 kar9Δ::KAN*$^{R}$ *GFP-TUB1::LEU2 ura3-52 lys2- 801 leu2-Δ1 his3-Δ200 trp1-Δ63* | YEF473 | This study |
| SMY1733 | Mata *dyn1*$^{R2543K}$*-3GFP::TRP1 NUP133-3mCherry::URA3 kar9Δ::KAN*$^{R}$ *GFP-TUB1::LEU2 ura3-52 lys2- 801 leu2-Δ1 his3-Δ200 trp1-Δ63* | YEF473 | This study |
| SMY1740 | Mata *DYN3-13MYC::HPH PAC11-13MYC::TRP ZZ-TEV-3HA-dyn1*$^{I554M}$*-HaloTag::KAN*$^{R}$ *nip100Δ pep4Δ::HIS5 prb1Δ his3-11,15 ura3-52 leu2-3,112 ade2-1 trp-1* | W303 | This study |
| SMY1744 | Mata *dyn1*$^{R241L}$*-3GFP::TRP1 NUP133-3mCherry::URA3 kar9Δ::KAN*$^{R}$ *GFP-TUB1::LEU2 ura3-52 lys2- 801 leu2-Δ1 his3-Δ200 trp1-Δ63* | YEF473 | This study |

| | | | |
|---|---|---|---|
| SMY1754 | Mata *DYN3-13MYC::HPH PAC11-13MYC::TRP ZZ-TEV-3HA-dyn1*$^{R2543K}$*-HaloTag::KAN*$^{R}$ *nip100Δ pep4Δ::HIS5 prb1Δ his3-11,15 ura3-52 leu2-3,112 ade2-1 trp-1* | W303 | This study |
| SMY1755 | Mata *DYN3-13MYC::HPH PAC11-13MYC::TRP ZZ-TEV-3HA-dyn1*$^{R2543K}$*-HaloTag::KAN*$^{R}$ *nip100Δ pep4Δ::HIS5 prb1Δ his3-11,15 ura3-52 leu2-3,112 ade2-1 trp-1* | W303 | This study |
| SMY1756 | Mata *DYN3-13MYC::HPH PAC11-13MYC::TRP ZZ-TEV-3HA-dyn1*$^{R3152N}$*-HaloTag::KAN*$^{R}$ *nip100Δ pep4Δ::HIS5 prb1Δ his3-11,15 ura3-52 leu2-3,112 ade2-1 trp-1* | W303 | This study |
| SMY1766 | Matα *dyn1*$^{K3160Q}$*-3GFP::TRP1 NUP133-3mCherry::URA3 kar9Δ::KAN*$^{R}$ *GFP-TUB1::LEU2 ura3-52 lys2- 801 leu2-Δ1 his3-Δ200 trp1-Δ63* | YEF473 | This study |
| SMY1774 | Mata *dyn1*$^{W612C}$*-3GFP::TRP1 NUP133-3mCherry::URA3 kar9Δ::KAN*$^{R}$ *GFP-TUB1::LEU2 ura3-52 lys2- 801 leu2-Δ1 his3-Δ200 trp1-Δ63* | YEF473 | This study |
| SMY1816 | Mata *dyn1*$^{H3639P,W3640G}$*-3GFP::TRP1 TUB1+3'UTR::HPH::HIS3p:mRuby2-TUB1 ura3-52 lys2-801 leu2-Δ1 his3-Δ200 trp1-Δ63* | YEF473 | This study |
| SMY1833 | Matα *dyn1*$^{L213I}$*-3GFP::TRP1 NUP133-3mCherry::URA3 kar9Δ::KAN*$^{R}$ *GFP-TUB1::LEU2 ura3-52 lys2- 801 leu2-Δ1 his3-Δ200 trp1-Δ63* | YEF473 | This study |
| SMY1834 | Mata *dyn1*$^{K1475Q}$*-3GFP::TRP1 NUP133-3mCherry::URA3 kar9Δ::KAN*$^{R}$ *GFP-TUB1::LEU2 ura3-52 lys2- 801 leu2-Δ1 his3-Δ200 trp1-Δ63* | YEF473 | This study |
| SMY1841 | Mata *dyn1*$^{F3638G,H3639P}$*-3GFP::TRP1 TUB1+3'UTR::HPH::HIS3p:mRuby2-TUB1 ura3-52 lys2-801 leu2-Δ1 his3-Δ200 trp1-Δ63* | YEF473 | This study |
| SMY1842 | Mata *dyn1*$^{F3638G,H3639P}$*-3GFP::TRP1 TUB1+3'UTR::HPH::HIS3p:mRuby2-TUB1 ura3-52 lys2-801 leu2-Δ1 his3-Δ200 trp1-Δ63* | YEF473 | This study |
| SMY1857 | Mata *dyn1*$^{R1822S,R1852V}$*-3GFP::TRP1 TUB1+3'UTR::HPH::HIS3p:mRuby2-TUB1 ura3-52 lys2-801 leu2-Δ1 his3-Δ200 trp1-Δ63* | YEF473 | This study |
| SMY1858 | Mata *dyn1*$^{R1852V}$*-3GFP::TRP1 TUB1+3'UTR::HPH::HIS3p:mRuby2-TUB1 ura3-52 lys2-801 leu2-Δ1 his3-Δ200 trp1-Δ63* | YEF473 | This study |
| SMY1866 | Mata *dyn1*$^{R1852V}$*-3GFP::TRP1 NUP133-3mCherry::URA3 kar9Δ::KAN*$^{R}$ *GFP-TUB1::LEU2 ura3-52 lys2- 801 leu2-Δ1 his3-Δ200 trp1-Δ63* | YEF473 | This study |

| | | | |
|---|---|---|---|
| SMY1867 | Matα *dyn1*$^{C1822S,R1852V}$*-3GFP::TRP1 NUP133-3mCherry::URA3 kar9Δ::KAN*$^{R}$ *GFP-TUB1::LEU2 ura3-52 lys2- 801 leu2-Δ1 his3-Δ200 trp1-Δ63* | YEF473 | This study |
| SMY1868 | Mata *DYN3-13MYC::HPH PAC11-13MYC::TRP ZZ-TEV-3HA-dyn1*$^{E109I}$*-HaloTag::KAN*$^{R}$ *nip100Δ pep4Δ::HIS5 prb1Δ his3-11,15 ura3-52 leu2-3,112 ade2-1 trp-1* | W303 | This study |
| SMY1883 | Mata *DYN3-13MYC::HPH PAC11-13MYC::TRP ZZ-TEV-3HA-dyn1*$^{N283R}$*-HaloTag::KAN*$^{R}$ *nip100Δ pep4Δ::HIS5 prb1Δ his3-11,15 ura3-52 leu2-3,112 ade2-1 trp-1* | W303 | This study |
| SMY1922 | Mata *dyn1*$^{F3638G,H3639P,W3640G}$*-3GFP::TRP1 TUB1+3'UTR::HPH::HIS3p:mRuby2-TUB1 ura3-52 lys2-801 leu2-Δ1 his3-Δ200 trp1-Δ63* | YEF473 | This study |
| SMY1923 | Mata *dyn1*$^{F3638G,H3639P,W3640G}$*-3GFP::TRP1 TUB1+3'UTR::HPH::HIS3p:mRuby2-TUB1 ura3-52 lys2-801 leu2-Δ1 his3-Δ200 trp1-Δ63* | YEF473 | This study |
| SMY1933 | Matα *dyn1*$^{F3638G,H3639P,F3640G}$*-3GFP::TRP1 NUP133-3mCherry::URA3 kar9Δ::KAN*$^{R}$ *GFP-TUB1::LEU2 ura3-52 lys2- 801 leu2-Δ1 his3-Δ200 trp1-Δ63* | YEF473 | This study |
| SMY1934 | Matα *dyn1*$^{F3638G,H3639P,F3640G}$*-3GFP::TRP1 NUP133-3mCherry::URA3 kar9Δ::KAN*$^{R}$ *GFP-TUB1::LEU2 ura3-52 lys2- 801 leu2-Δ1 his3-Δ200 trp1-Δ63* | YEF473 | This study |
| SMY1959 | Mata *DYN3-13MYC::HPH PAC11-13MYC::TRP ZZ-TEV-3HA-dyn1*$^{F3638G,H3639P,W3640G}$*-HaloTag::KAN*$^{R}$ *nip100Δ pep4Δ::HIS5 prb1Δ his3-11,15 ura3-52 leu2-3,112 ade2-1 trp-1* | W303 | This study |
| SMY1960 | Mata *DYN3-13MYC::HPH PAC11-13MYC::TRP ZZ-TEV-3HA- dyn1*$^{F3638G,H3639P,W3640G}$*-HaloTag::KAN*$^{R}$ *nip100Δ pep4Δ::HIS5 prb1Δ his3-11,15 ura3-52 leu2-3,112 ade2-1 trp-1* | W303 | This study |
| SMY2104 | Mata *DYN3-13MYC::HPH PAC11-13MYC::TRP ZZ-TEV-3HA-dyn1*$^{R1852C}$*-HaloTag::KAN*$^{R}$ *nip100Δ pep4Δ::HIS5 prb1Δ his3-11,15 ura3-52 leu2-3,112 ade2-1 trp-1* | W303 | This study |
| SMY2129 | Matα *dyn1*$^{C1822A,R1852C}$*-3GFP::TRP1 NUP133-3mCherry::URA3 kar9Δ::KAN*$^{R}$ *GFP-TUB1::LEU2 ura3-52 lys2- 801 leu2-Δ1 his3-Δ200 trp1-Δ63* | YEF473 | This study |
| SMY2158 | Matα *DYN1-3GFP::TRP1 TUB1+3'UTR::HPH::HIS3p:mRuby2-TUB1 PDR1::pdr1-DBD-CYC8::LEU2 ura3-52 lys2- 801 leu2-Δ1 his3-Δ200 trp1-Δ63* | YEF473 | This study |

| | | | |
|---|---|---|---|
| SMY2162 | Mata *dyn1*$^{H3639P}$*-3GFP::TRP1 TUB1+3'UTR::HPH::HIS3p:mRuby2-TUB1 PDR1::pdr1-DBD-CYC8::LEU2 ura3-52 lys2- 801 leu2-Δ1 his3-Δ200 trp1-Δ63* | YEF473 | This study |

#### SUPPLEMENTAL REFERENCES


1. Schmidt, H., Zalyte, R., Urnavicius, L. & Carter, A. P. Structure of human cytoplasmic dynein-2 primed for its power stroke. *Nature* 518, 435-438, (2015).
2. Redwine, W. B. *et al.* Structural basis for microtubule binding and release by dynein. *Science* 337, 1532-1536, (2012).
3. Urnavicius, L. *et al.* Cryo-EM shows how dynactin recruits two dyneins for faster movement. *Nature* 554, 202-206, (2018).
4. Fraley, C. & Raftery, A. E. Bayesian Regularization for Normal Mixture Estimation and Model-Based Clustering *Journal of Classification* 24, 155-181, (2007).
5. Ecklund, K. H. *et al.* She1 affects dynein through direct interactions with the microtubule and the dynein microtubule-binding domain. *Nature communications* 8, 2151, (2017).
